# Non-Canonical Ago Loading of EV-Derived Exogenous miRNA Generates Foreign miRNPs on Endosomes to Arbitrate Gene Expression in Recipient Cells

**DOI:** 10.1101/2020.05.26.115899

**Authors:** Bartika Ghoshal, Edouard Bertrand, Suvendra N. Bhattacharyya

**Affiliations:** RNA Biology Research Laboratory, Molecular Genetics Division, CSIR-Indian Institute of Chemical Biology; Institut de Génétique Moléculaire de Montpellier, Université de Montpellier, CNRS, Montpellier, France

**Keywords:** Non-canonical miRNA loading of Ago, Extracellular Vesicles-mediated miRNA transfer, miRNA endocytosis, miRNA-activity regulation by endosomes, Dicer1-independent miRNA loading, Ago and edosomes

## Abstract

MicroRNAs, the tiny regulators of gene expression, can be transferred between neighbouring cells via Extracellular Vesicles (EV) to control the expression of genes in both donor and recipient cells. How the Extracellular Vesicles-derived miRNAs are getting internalized and become functional in target cells is an unresolved question. We found that EV-associated miRNAs are primarily single stranded and, to become functional, get loaded with the Ago proteins present in the recipient cells without requiring host Dicer1. Following endocytosis of miRNA-containing extracellular vesicles, loading of host Ago occurs on the endosomal membrane where pH-dependent membrane fusion triggers the release of internalized miRNAs to form exogenous miRNP pool. In addition, loading of exogenous miRNA to host Ago also depends on the mitochondrial activity of recipient cells. Internalization of hepatocyte derived miR-122 containing EVs in macrophage activates the recipient cell by targeting cytokine expression. *Leishmania donovani*, a protozoan parasite known to affect endocytosis and to cause mitochondrial depolarization in infected macrophages, restricts the EV-internalization process and thereby preventing inflammatory cytokine expression and ensuring internalized pathogen survival in macrophage.

## Introduction

Intercellular communication in metazoan cells within the same or different tissues can be achieved via exchange of membrane enclosed vesicles that can carry proteins or nucleic acids (Maas et al, 2017). This exchange of materials ensures a robust homeostasis in metabolic processes in animal cells and thus considered as a very important physiological phenomenon in animals (Stahl & Raposo, 2019). These extracellular vesicles (EVs), as they are collectively known as, either originate from the multivesicular bodies (MVBs) or the plasma membrane and are classified according to their origin and physical status as microvesicles, ectosomes, microparticles or exosomes (van Niel et al, 2018). Exosomes the 40-100nm sized vesicles, positive for the marker protein CD63, and are formed during membrane invagination of late endosomes to generate multivesicular bodies (MVBs) followed by MVB fusion to cell membrane (Colombo et al, 2014). Extracellular vesicles or exosomes, in particular, have been detected in diverse body fluids (Raposo & Stoorvogel, 2013), and are known to carry different cargoes like mRNAs, microRNAs, proteins, macromolecules to the recipient cell to ensure an intercellular communication and exchange of materials. MicroRNAs or miRNAs are 20-22 nucleotide long gene regulatory RNAs which by base pairing to target messages can trigger translational repression or degradation of target RNAs (Fabian et al, 2010; Filipowicz et al, 2008). miRNAs target important regulatory genes, and have been found to control growth of cancer cells and to play crucial role in pathogenesis of other diseases as well. miRNAs have been found to be present in the body fluids either as ‘free’ entries (sometimes in complex with the Argonaute proteins) or in enclosed vesicles like exosomes (Arroyo et al, 2011; Kosaka et al, 2010).

Transfer of functional microRNAs by exosomes or extracellular vesicles (EVs) to recipient cells has been observed to evoke a physiological response both in cancer or immune cells (Kogure et al, 2011; Mittelbrunn et al, 2011; Montecalvo et al, 2012; Pegtel et al, 2010; Salido-Guadarrama et al, 2014; Valadi et al, 2007). EV-mediated crosstalk between cells having different miRNA expression profiles has also been reported (Basu & Bhattacharyya, 2014). MicroRNAs are sorted selectively for packaging into Multivesicular bodies(MVBs) by specific sets of proteins before the MVBs can fuse with cell membrane to release the EVs, packed with specific miRNAs in the intercellular space in a context dependent manner (Cha et al, 2015; Mukherjee et al, 2016; Shurtleff et al, 2016; Villarroya-Beltri et al, 2013). Different miRNA associated proteins like the Ago2 and components of the RISC loading complex have also been found to be involved in exosomal export of miRNAs in different cells (Lv et al, 2014; Melo et al, 2014).

On investigating the internalization process of EVs into recipient cells, it has been found to occur either by endocytosis (Svensson et al, 2013; Tian et al, 2010), macropinocytosis (Tian et al, 2014), phagocytosis (Feng et al, 2010) or engulfment with the help of filopodia (Heusermann et al, 2016). However, despite recent advances in the study of EV-mediated transport of miRNA, the mode of uptake of exosomal miRNA and the factors that render them functional in recipient cells are entirely unexplored. Using an ectopic miRNA expression system in human HeLa cells, we have observed functional miRNA transfer to the recipient cell by EVs. This exogenous miRNA enters the recipient cell in a single-stranded form and in a Dicer independent manner. Internalized miRNAs become associated with the recipient cell Argonaute proteins. We have observed that the internalized miRNAs utilise the endocytic pathway to reach the Endoplasmic Reticulum (ER) of the target cells to elicit their repressive action. We have substantiated our data in an *in vitro* assay to show that the decrease of endosomal pH causes the release of the miRNA from the internalized EVs – a prerequisite process for functional loading of internalized miRNAs to recipient cells’ Ago2. In this context, we have observed that the miR-122, a hepatic miRNA, can up regulate pro-inflammatory cytokines in recipient macrophage cells following its entry. Conversely upon infection *by Leishmania donovani (Ld),* a protozoan parasite and the causative agent of visceral leishmaniasis in humans, we have found that the resident parasite restricts the EV-mediated internalization of miR-122 and thereby prevents up regulation of the pro-inflammatory cytokines for sustained infection in the recipient macrophages. EV-entry was found to be restricted by mitochondrial depolarization happening in *Ld* infected cells also defective in endosomal pathways essential for EV-mediated miRNA entry.

## RESULTS

### Functional miRNA transfer between mammalian cells via EVs

To understand the mechanistic aspect of uptake of EV enclosed miRNAs, we used engineered HeLa cells ectopically expressing a “foreign” liver specific miR-122 as donor cells that should produce EVs having miR-122 packed into it. The miR-122 is expressed abundantly in the liver cells that are otherwise not detectable in HeLa cells (Lagos-Quintana et al, 2002; Landgraf et al, 2007). Non-engineered HeLa cells without miR-122 expression cassette thus could serve as recipient cells where EV-mediated uptake and functional activity of transferred miR-122 can be measured. Increase of miR-122 in recipient HeLa cells, is the measure of the miRNA-transferred after the pre-treatment of HeLa cells with EVs derived from miR-122 expressing HeLa cells (Figure 1A).

**Figure 1.**
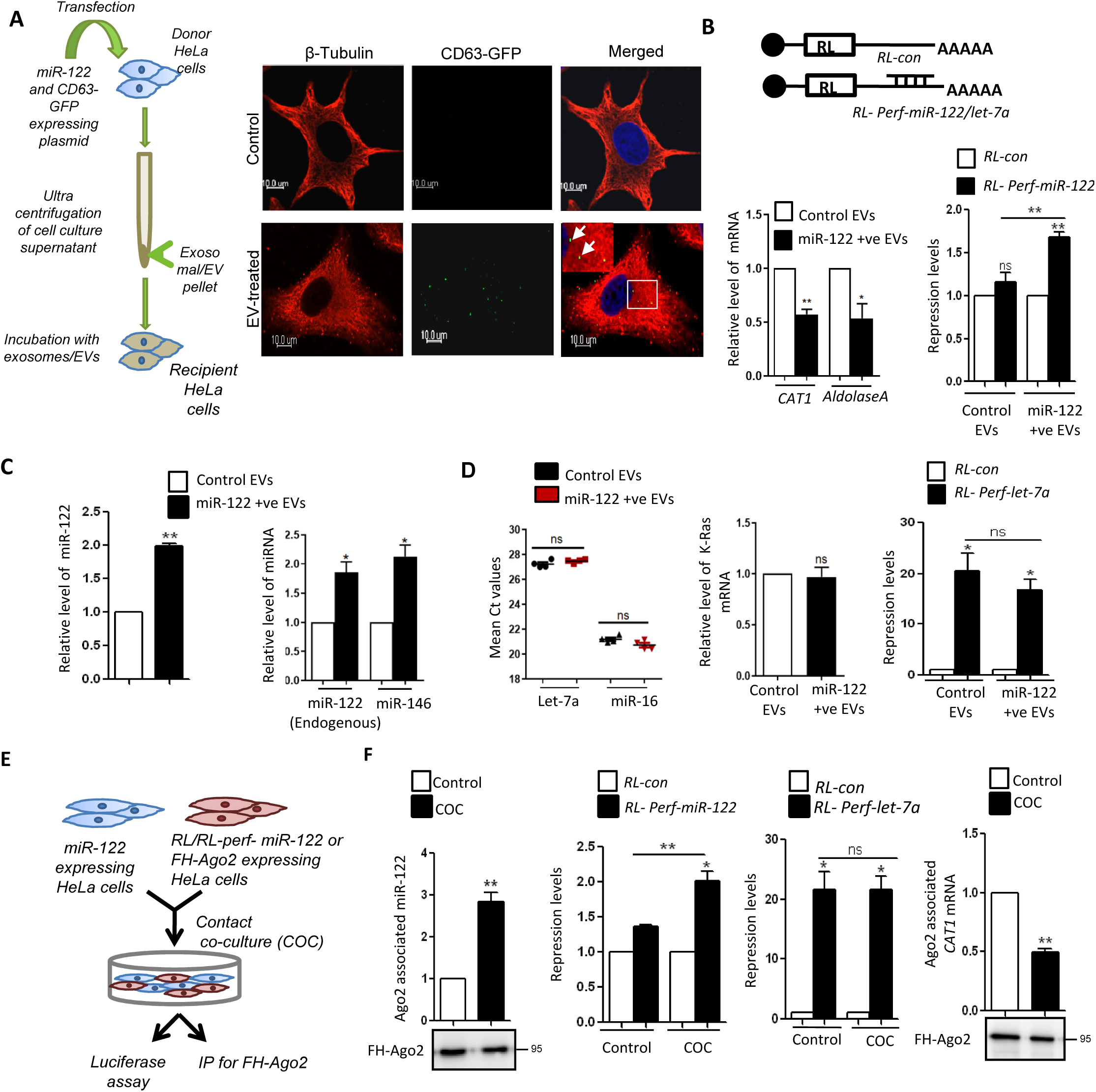
EV-mediated functional miRNA transfer across cell boundaries in heterogeneous context. (A) Schematic representation of EV-mediated miR-122 transfer in non-hepatic HeLa cells. EVs isolated by ultracentrifugation from miR-122 expressing HeLa cells were used for treatment of miR-122 negative HeLa recipient cells. In the Right panel, confocal image is used to show internalization of EVs in recipient HeLa cells following incubation with EVs isolated from CD63-GFP expressing HeLa cells. The cell boundary denoted in red is for β-tubulin network detected simultaneously while imaging was done for GFP-CD63. The green spots marked by white arrows are internalized CD63-GFP vesicles. Scale bar 10 µm. Inset shows zoomed part of a the image showing internalized EVs marked by white arrows. (B) Activity analysis of EV-transferred miR-122 in HeLa cells. The *top* panel shows the scheme of the reporter mRNA used to study the repression activity of EV-transferred miR-122 in recipient cells. Relative levels of *CAT1, ALDOLASE* mRNAs, known endogenous miR-122 targets, in recipient HeLa cells treated or untreated with miR-122 containing EVs (mean ± S.E.M., n=3). Relative repression level of a *Renilla*-based-miR122 reporter with one perfect miR-122 binding site in recipient cells treated or untreated with miR-122 containing EVs has also been shown in right *bottom* panel. Expression from firefly luciferase reporter co-transfected with RL reporter was used as transfection control and value normalization (mean ± S.E.M., n=3). (C) Internalization of single stranded mature miR-122 in HeLa cells following miR-122 containing EV treatment. Real Time PCR analysis was done to estimate the internalization of Ev-derived miR-122 in recipient HeLa cells (*left panel*). U6 snRNA was used as a normalizing control (mean ± S.E.M., n=3). Level of internalization of exsomal miR-146a when expressed in donor HeLa cells via EVs to recipient HeLa. The value was compared against miR-122 level also expressed in donor HeLa cells (right panel) (mean ± S.E.M., n=3). (D) No change in endogeneous miRNA content or activity upon treatment of HeLa cells with miR-122 containing EVs. Real Time PCR was used to show no difference in the Ct values of endogenous miRNA let-7a and miR-16 in cells untreated or treated with miR-122 containing exosomes (n=4) (*left panel*). Relative level of K-Ras, an endogenous target of let-7a, in recipient cells treated or untreated with miR-122 positive EVs were analyzed (*middle panel*). Relative repression level of a *Renilla*-based-let7a reporter with a perfect let-7a binding site in recipient cells treated or untreated with miR-122 postive EVs (*right panel*). (E) Schematic representation of contact co-culture model of EV-mediated miR-122 transfer in HeLa cells. Donor HeLa cells were transfected with miR-122 expression plasmid and co-cultured with the recipient HeLa cells expressing FLAG-HA-Ago2 or luciferase reporters. (F) Transfer of miR-122 repressive activity between HeLa cells and functional transfer of mature miR-122 in recipient cell Ago2 protein. In the *left* panel, Ago2 associated miR-122 levels in recipient cells co-cultured with or without miR-122 expressing HeLa cells. The middle panel shows repression levels of miR-122 and let-7a reporters in HeLa cells co-cultured with HeLa cells expressing or not expressing miR-122. In the *right* most panel, Ago2 associated *CAT1* mRNA levels in recipient Hela cells co-cultured with donor HeLa cells with or without miR-122 expression. The mRNA and miRNA levels were normalized with the band intensity of immunoprecipitated FLAG-HA-Ago2. For statistical significance, minimum three independent experiments were considered unless otherwise mentioned and error bars are represented as ± S.E.M. P-values were calculated by utilising Student’s t-test. ns: non-significant, *P < 0.05, **P < 0.01, ***P < 0.0001.

The isolated EVs from donor HeLa cells expressing miR-122 were subjected to atomic force microscopy and nano particle tracking analysis (NTA) to get the measurement of their shapes and sizes and were compared to EVs derived from HeLa cells not expressing miR-122 (Figure S1A and D). No difference in sizes, concentrations as well as in protein content was observed between control and miR-122 containing EVs (Figure S1B, S1C). For microscopic analysis, the donor cells were also transfected with CD63-GFP, which were expressing GFP tagged exosomal marker protein tetraspanin CD63. The EVs with CD63-GFP were collected and were added to recipient cells. 3D imaging on recipient cells showed internalization of the GFP-tagged CD63 containing EVs (Figure 1A).

We then further confirmed the presence of mature miR-122 in the EVs isolated from miR-122 expressing donor HeLa cell. The levels of mature miR-122 detected in EVs from donor HeLa cells remained unchanged upon RNase-free DNase treatment of the nucleic acids recovered from miR-122 positive EVs. This data has ruled out the possible presence of miR-122 encoding DNA fragment as contaminant in the EVs used for the treatment of recipient HeLa cells to cause increased miR-122 detected there upon EV treatment (Figure S1E). The miR-122* was also not detected in recipient HeLa and was in the non-reliable detection limit in EVs derived from miR-122 expressing HeLa cells. The average copy number of mature miR-122 per EVs was 1.95 while the total amount of miR-122 transferred to recipient HeLa after treatment for 16h was just above 10,000 molecules per cells. No pre-miR-122 was detected in recipient HeLa cells treated with mature miR-122 positive EVs.

Upon transfer of miR-122 via EVs, we found decreased expression of miR-122 target genes in EV-treated cells that signifies functional fitness of the EV-mediated transferred miR-122 across donor and recipient HeLa cell boundaries. The transferred miR-122 was found to cause downregulation the target messages (Figure 1B). The mature miR-122 content was found to increase in the recipient cells treated with miR-122 containing EVs (Figure 1C). The endogenous let-7a and miR-16 miRNA levels as well as their repressive activities remained unchanged between the EV-treated groups of recipient cells (Figure 1D). Similar to miR-122, miR-146 also could get transferred to recipient HeLa cells via EVs when ectopically expressed in donor HeLa cell (Figure 1C).

One possible mechanism that ensures miRNA homeostasis in animal cells could be the EV-mediated transfer of miRNAs between neighbouring cells. This may also happen between cells in contact co-culture. In order to revalidate the exosomal uptake of miRNA in recipient HeLa cells, when the donor and recipient cells are in contact, we adopted a co-culture technique to follow the miRNA transfer across cellular boundaries (Figure 1E). This technique has been used earlier to deduce the functional transfer of miRNA between cells (Basu & Bhattacharyya, 2014). The data was consistent with the miRNA uptake observed in HeLa cells treated with miR-122 containing EVs. By co-culturing two different pools of HeLa cells; one expressing miR-122 and other with a miR-122 reporter, we can test functionally the activity of transferred miRNA across cell boundaries (Figure 1F). We noted transfer of repressive activity of miR-122 as expected in cells having differential expression of miR-122 and that was without having any effect on endogeneous let-7a miRNA activity.

### The internalized miRNAs associates with Ago2 in recipient cells in a Dicer-independent manner

Having standardised the model systems for EV uptake studies, we followed the uptake process over time. By incubating recipient cells with EVs derived from donor cells expressing GFP-tagged CD63, we estimated the uptake by following the CD63-GFP internalization in recipient cells over time. The recipient cells were incubated with CD63-GFP positive EVs at different time points of 0, 4 and 16 hours and internalization of GFP-CD63 was quantified microscopically. The internalization of CD63-GFP increased with time till 16h of observation (Figure 2A,B). We also measured the internalized miR-122 level and its incorporation with Ago2 of the host cells to document a time dependent increase (Figure 2C). Interestingly, incubation beyond 16h showed a decreased miR-122 uptake (data not shown). Transfer of internalized miRNAs to recipient cell Ago2 was also documented in contact co-culture model (Figure 1F). Transfer of EV-derived miRNAs to Ago proteins was found to be not exclusive for Ago2 only but Ago3 was also found to be loaded with internalized miRNAs while low level of miRNA was found to be bound to control NHA-LacZ protein when expressed in recipient HeLa cells and was used as negative control (Figure 2D).

**Figure 2.**
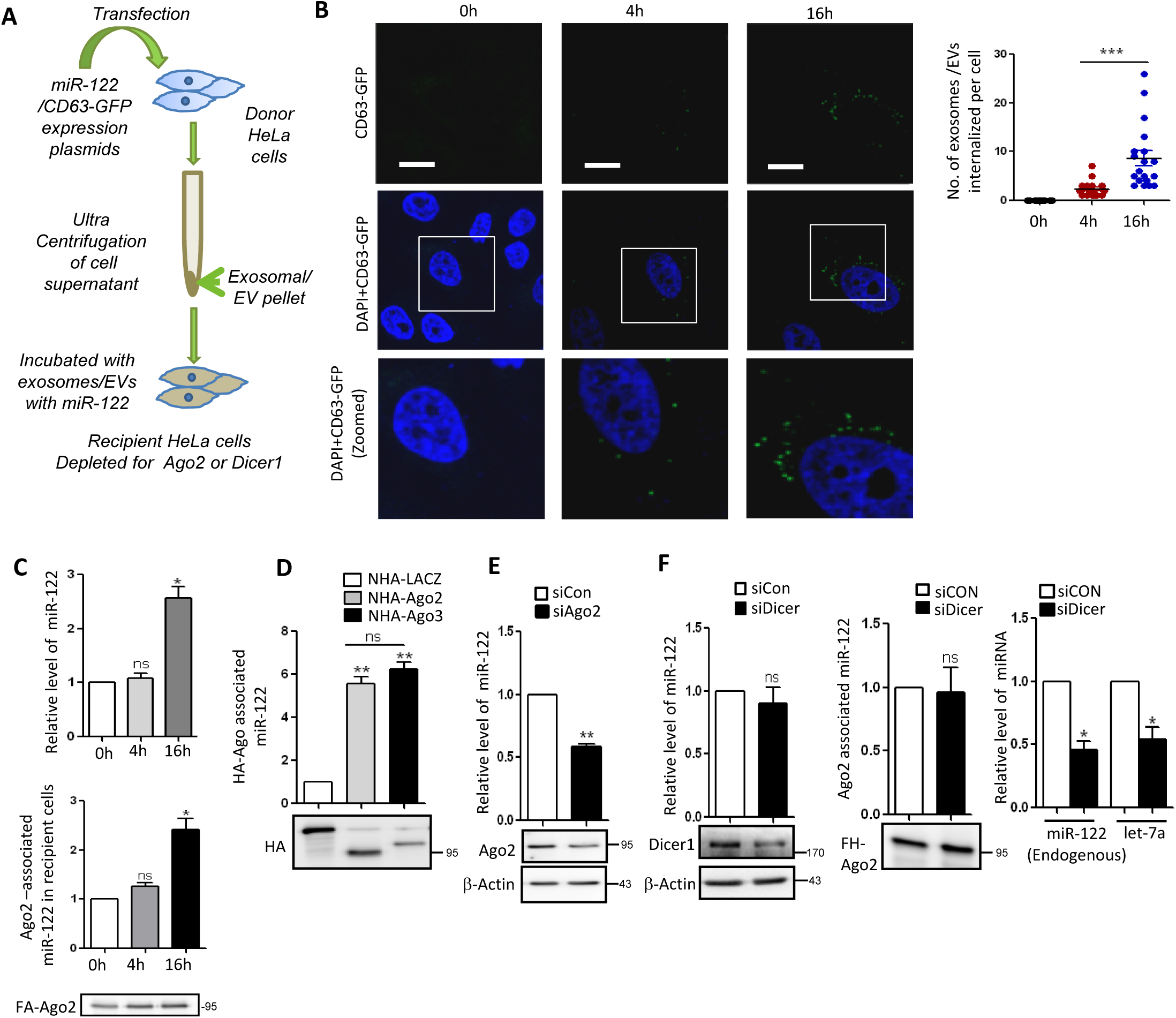
Internalization and Dicer independent Ago2 loading of EV-derived miRNAs in recipient cells. (A) Schematic representation of the assay to study the transfer of EV-derived miRNAs to recipient cell Ago2. Using HeLa cells knocked down for Ago2 or Dicer were treated with EVs derived from another batch of HeLa cells ectopically expressing miR-122. For microscopic analysis, the donor cells are co-transfected with CD63-GFP encoding plasmid followed by isolation of CD63-GFP tagged EVs. These are used for treatment of recipient cells and imaging was done for internalization of CD63-GFP positive EVs in recipient HeLa cells at different time points of 0, 4, 16 hours. (B) Imaging of recipient cells after 0, 4 and 16 hours of CD63-GFP positive EV-treatment. Detection of internalization of CD63-GFP EVs was monitored microscopically. Graphical representation of the quantification done for number of EV internalized at different time points (n=19) (top *right panel*). (C) Time dependent increase of cellular (*upper panel)* and Ago2 associated (*lower panel*) miR-122 in recipient cells. Relative levels of miR-122 were measured in recipient cells at different time points of treatment at 0, 4, 16 hours with EVs packed with miR-122. U6snRNA was used as a normalizing control. The panel depicts the Ago2 associated internalized miR-122 levels in recipient cells at the aforementioned time points. Amount of Ago2 immunoprecipitated were used for normalization of miR-122 content. (D) Association of EV-derived miR-122 with recipient cell Ago2 or Ago3 in contact co-culture model to study EV-mediated miRNA transfer across cell memebrane. Associated miRNA values were normalized to immunoprecipitated levels of NHA-Ago2 or Ago3 levels (lower panel). NHA-LacZ was used as a negative control. (E) Effect of depletion of Ago2 on relative level of miR-122 internalization in recipient cells. Levels of miR-122 were measured in HeLa cells transfected with siCon or siAgo2 and treated with miR-122 containing EVs. U6 was used for normalization. Western Blot was used to show knockdown of Ago2. β-Actin was used as loading control. (F) Effect of depletion of Dicer1 on levels and Ago2 association of EV-derived miR-122 in HeLa cells expressing FH-Ago2. Relative level of total miR-122 internalized after 16h of treatment of recipient HeLa transfected with siCon or siDicer with EVs positive for miR-122 (mean ± S.E.M., n=4). Western blot was done to show knockdown of Dicer. β-Actin was used as loading control (*left panel*). Ago2 associated miR-122 was also measured in recipient cells transfected with siCon or siDicer in contact co-culture model (*middle panel*). Relative levels of endogenous let-7a and miR-122 (when expressed as pre-miR-122) in HeLa cells co-transfected with miR-122 expressing plasmid and siCon or siDicer (*right panel*). For statistical significance, minimum three independent experiments were considered in each case unless otherwise mentioned and error bars are represented as mean ± S.E.M. P-values were calculated by utilising Student’s t-test. ns: non-significant, *P < 0.05, **P < 0.01, ***P < 0.0001.

In the canonical pathway, Ago2 loading of miRNA is dependent on its interaction with Dicer1 and processing of pre-miRNA by Dicer1 to generate the double stranded miRNAs. The miRNA sense strand then gets loaded to Ago proteins in a Dicer1 dependent manner (Frohn et al, 2012). To test whether the internalization and Ago-association of internalized single stranded miRNA is dependent on the prime components of the miRNA biogenesis pathway in the recipient cell, Ago2 and Dicer1 proteins of the recipient cell were targeted by specific siRNAs. When the recipient cells were knocked down for Ago2, the internalization of EV-transferred miR-122 decreased in the recipient cell suggesting a requirement of recipient cell Ago2 for the internalization and function of the EV-transferred miR-122 (Figure 2E).

When the recipient cells were knocked down for Dicer1 and incubated with miR-122 containing EVs, the amount of internalized miRNAs remained unchanged as compared to the control. Ago2 associated miR-122 levels in the control and Dicer1 knockdown cells also remained unchanged (Figure 2F). However, in HeLa cells co-transfected with pre-miR-122 expression plasmid and siDicer, the levels of mature miR-122 was reduced significantly along with a similar decrease, as expected, for endogenous let-7a levels upon Dicer1 knockdown (Figure 2F). Therefore, being independent of Dicer1 presence in recipient cells, the data on internalization of EV-derived miRNA and its Ago-loading suggests that the transfer of miR-122 via EVs involves the single-stranded and mature form of the miRNA (Jeppesen et al, 2019). If the precursor miRNA (pre-miR-122) would have been transferred then miRNA incorporation to recipient cell Ago2 was expected to be affected upon knockdown of Dicer1 as Dicer1 is required for pre-miRNA processing. Dicer1 independent non-canonical Ago-loading of EV derived mature miRNA is possibly required to elicit a quick and effective response in the recipient cells.

### EV-derived miRNAs enters recipient cell via the endosomal pathway

To understand how the EV-derived miRNA enters the recipient cell and where does it get finally localized, the involvement of specific proteins known to control endocytic processes were targeted. Dynamin2 is needed for pinching off the vesicles into the interior of the cell from cell membrane and is required for the internalization process in certain endocytic pathways (De Camilli et al, 1995; Feng et al, 2010). Upon knockdown of Dynamin2 in the recipient cell, the internalization of miRNA was significantly reduced. The dependence of EV-internalization on Dynamin2 was also confirmed in microscopic analysis of CD63-GFP tagged vesicles internalization in recipient HeLa cells where Dynamin 2 expression was reduced by siRNA (Figure S2A, S2B). This suggests that Dynamin2, a protein prerequisite for endocytosis is also needed for the internalization of EV-containing miRNAs.

Since Dynamin2 has been found to be involved in the uptake pathway of the EV-derived miRNA, we wanted to assess the sub-cellular localization of the internalized miRNA in the recipient cells. Cellular lysates of recipient cells post 16h EV-treatment was collected in isotonic condition and were analyzed on Optiprep^R^ density gradient to separate the organelles. Western blot analysis was done with different fractions to demarcate different sub-cellular compartments (Figure 3A, 3B). The different sub-cellular fractions were collected and pooled together depending on the presence of marker proteins for different organelles and RNA was isolated from the different pooled fractions. Fractions 2, 3 were enriched for early endosomes while fractions 4-6 represented late endosomes and lysosomes and fractions 7-9 had enrichment for ER marker protein. In a steady state context, the majority of the internalized EV-derived miRNAs (total and Ago2 associated) were found to be associated with ER (Fig 3C). Thapsigargin is an ER-stress inducer and treatment of recipient cells with Thapsigargin followed by incubation with miR-122 containing EVs, resulted in defective internalization of EV-derived miRNA-122 in treated cells (Figure S2C, S2D). Therefore fitness of ER controls the internalization and functioning of the EV-derived miR-122 in human cells. Unlike the Dicer1 dependent canonical pathway where availability of the target messages has a positive influence of cognate miRNA formation (Bose & Bhattacharyya, 2016), we detected no increase of EV-delivered miRNA content in presence of its target messages in recipient cells, rather a decrease in total miRNA content was noted in addition to no change in Ago2-associated miRNA content was detected (Figure S2F, S2G). This data further supports a non-canonical miRNA loading pathway prevalent for EV-internalized miRNAs.

**Figure 3.**
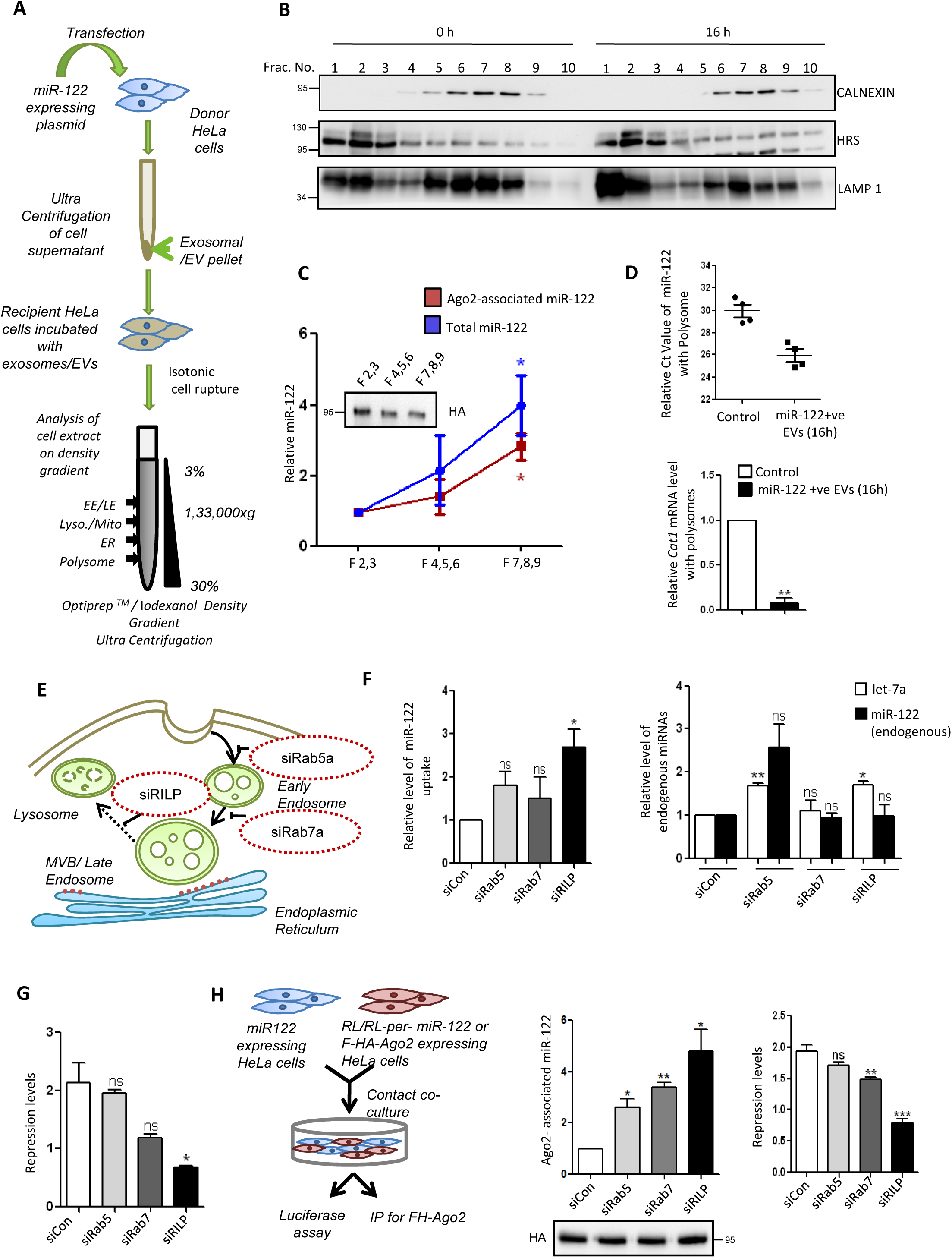
The EV-transferred miRNA reaches the Endoplasmic Reticulum of the recipient cell by utilising the endocytic pathway. (A) Schematic representation of the sub-cellular fractionation of the EV-treated recipient cells to follow the localisation of EV-transferred miRNA. The recipient cells were incubated with miR-122 containing EVs followed by lysis and 3-30% Optiprep density gradient ultra-centrifugation to separate the different sub-cellular fractions. Individual fractions enriched for endosomal or ER marker proteins were pooled separately and RNA isolated from these fractions were quantified by qRT-PCR. (B) Western blot analysis of different marker proteins to demarcate the different subcellular fractions enriched for either Endoplasmic Reticulum (Calnexin), early endosomes (HRS), lysosomes (Lamp1) by density gradient ultra-centrifugation. (C) Optiprep density gradient ultracentrifugation of lysates from EV-treated recipient cells followed by quantitative Real Time PCR estimation of miR-122 in pooled fractions. Immunoprecipitation of FH-Ago2 from pooled fractions was done to see the subcellular location and Ago2 association of internalized EV-derived miR-122 (n=4). Ago2 levels in pooled fractions were estimated in accompanying western blot and used for normalization of Ago2 associated miRNA levels. (D) Polysome association of transferred miR-122 in recipient cells. Ct values of transferred miR-122 associated with polysomes in recipient HeLa cells. Values in untreated cells was used as control (n=4) (upper panel). The lower panel shows the endogenous miR-122 target *CAT1* mRNA levels in the polysomes of recipient cells after miR-122 containing EV treatment. (E) Schematic representation of the endocytic pathway in human cells. Some of the key proteins like Rab5a, Rab7a and Rab Interacting Lysosomal Protein (RILP) which are involved in different steps of endosome maturation pathway were knocked down individually by treating recipient HeLa cells with respective siRNAs to check their effect on pathway of internalization of EV-derived miRNAs. (F) Relative level of internalized EV-derived miR-122 in recipient cells transfected with siCon, siRab5, siRab7 or siRILP (*left panel*). Relative level of miR-122 and endogenous let-7a levels in cells co-transfected with miR-122 expressing plasmid and either with siCon, siRab5, siRab7 or siRILP to show difference in endogenous level of miRNA (let-7a) as compared to internalized miRNA (miR-122) via EVs (*right panel*). (G) Repression levels of RL-perfmiR-122 a miR-122 reporter plasmid in cells co-transfected with siRNAs against the endocytic pathway components and treated with miR-122 containing EVs. (H) Effect of depletion of endocytic pathway proteins on internalization of EV-derived miRNA in contact co-culture model of donor and recipient cells (*left panel*) was followed by quantifying the Ago2 associated miRNA levels (*middle panel*) and repression levels for miR-122 reporter RL-perfmiR-122 (*right panel*). For statistical significance, minimum three independent experiments were considered unless otherwise mentioned and error bars are represented as mean ± S.E.M. P-values were calculated by utilising Student’s t-test. ns: non-significant, *P < 0.05, **P < 0.01, ***P < 0.0001.

It has been shown earlier that ER membrane-located polysomes are the sites of miRNA-target RNA interaction (Barman & Bhattacharyya, 2015). Therefore it was expected to find the functional miRNPs with the ER attached polysomes even for the EV-transferred miRNAs. We wanted to explore the same by looking at the level of internalized miRNAs localized with polysomes. From the relative Ct value analysis it seems that there has been increase in polysomal miR-122 content upon treatment of recipient cells with miR-122 containing EVs. This was accompanied by a concomitant decrease in the target CAT1 mRNA level with the polysome (Figure 3D).

To understand how the internalized miRNA-containing EVs release their content to functionally transfer the miRNA to the recipient cell Ago2 for effective repressive activity, we explored the effect of depletion of specific protein components of the endosomal maturation pathways in mammalian cells to score the effect on EV-mediated miRNA entry and its function. The endosomal pathway comprises of distinct membrane bound compartments, which internalize molecules from the plasma membrane and recycle them back to the surface or sort them for lysosomal degradation. Three proteins of the endosomal pathway, Rab5a (an early endosomal GTPase required for early to late endosomal maturation), Rab7a (a late endosomal GTPase required for endosome maturation and lysosome interaction) and RILP (Rab Interacting Lysosomal Protein which facilitate late endosomes interaction with lysosomes) were targeted specifically to dissect the subcellular steps of endocytosis required for functional miRNA transfer to ER attached polysomes (Figure 3E). In a steady state, endosome numbers were not found to significantly alter on knockdown of the endosomal proteins individually (Figure S3A, S3B). Measuring the effect of knockdown of these factors, we found a stronger effect of RILP depletion on internalized miRNA content. However, the knockdown of RILP marginally altered the levels of endogenous let-7a or exogenously expressed miR-122 in recipient HeLa cells (Figure 3F). Interestingly Rab5 depletion had similar effect on both EV-internalized miR-122 and endogenous miRNAs and an increase in overall miRNA content in each case was detected. Surprisingly, the increase in miR-122-content did not get reflected in the repressive activity of the miR-122. Rather a reduction in miRNA-repressive activity was noted after knockdown of Rab5 or Rab7 and more significantly upon RILP depletion (Figure 3G). Using the co-culture model of miRNA-transfer, we did observe an increase in Ago2-incorporation of internalized miRNAs with depletion of Rabs and RILP but with a decrease in repressive activity (Figure 3H).

The increased miRNA levels with Ago2 upon Rab7 and RILP depletion with a decrease in miRNA repressive activity suggests that the newly formed miRNP does not reach its target, perhaps because of an altered localization when endosomal pathways are altered.

### Late endosomal maturation and low pH is a requisite for membrane fusion of endocytosed EV and Ago2 loading of EV-derived miRNAs

From the analysis described in the above experiments, it seems that majority of the internalized miRNAs get localized with ER-attached polysome at steady state upon treatment of HeLa cells with miR-122 loaded EVs. With depletion of endogenous Rab7 protein and RILP individually, we observed an increase in miRNA content in HeLa cells and the increased miRNA became Ago2 associated but got specifically enriched with endosome fraction (Figure 4A-B). This data suggest that the EV-derived miRNA is released in the host cell at the levels of endosomes, and a simple possibility would be that this occurs by fusion of the endocytosed EV membrane with the surrounding endosomal membrane.

**Figure 4.**
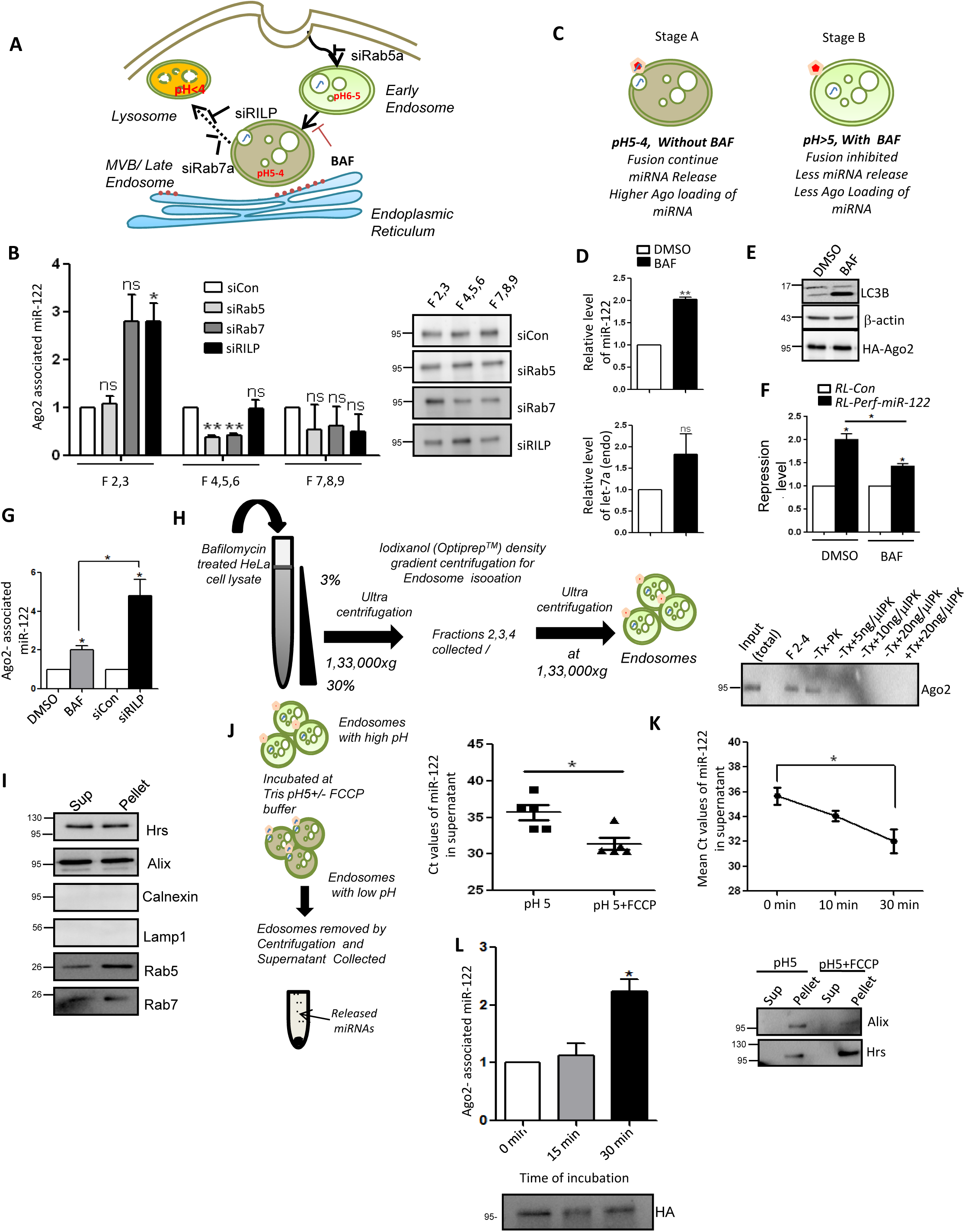
The release of internalized EV located miRNAs is dependent on pH change in endosomal compartment. (A) Diagrammatic depiction of the endocytic pathway shows the gradual change in pH reported to happen in each of the compartment during endosomal maturation. (B) Presence of Ago2 associated internalized miRNAs with endosomal and ER fractions in recipient cells. Optiprep density gradient ultracentrifugation of cell lysates from siCon, siRab5, siRab7 or siRILP transfected recipient cells after co-culture with cells expressing miR-122, were used to separate organelles based on their densities and fractions positive for Endosomal (2-3), lysosomal(4-6) and ER(7-9) marker proteins were pooled together followed by immunoprecipitation of FH-Ago2 from these pooled fractions. miRNAs associated with Ago2 in individual fractions were measured. (C) A schematic presentation of effect of Bafilomycin, an inhibitor of V-type ATPase, on EV-derived miRNA loading on recipient cell Ago2. Without Bafilomycin that otherwise prevent acidification of endosomal compartment, the miRNA containing internalized EVs fuses with endosomal membrane and miRNA should get loaded to Ago2 located on endosmal membrane. This effect is reduced on treatment of cells with Bafilomycin. (D) Relative level of EV-derived miR-122 in recipient HeLa cells treated with 25 nM Bafilomycin. DMSO was added to control cells (upper panel). Lower panel shows level of endogenous let-7a in recipient HeLa cells treated or untreated with Bafilomycin. (E) Western Blot of LC3B to show the increased cleavage of LC3B in cells treated or untreated with Bafilomycin. (F) Repression levels of RL-perfmiR-122, a miR-122 reporter in cells treated with miR-122 containing EVs and DMSO or Bafilomycin. (G) Relative levels of Ago2 associated EV-derived miR-122 in recipient cells. miRNA-associated with Ago2 immunoprecipitated from DMSO or Bafilomycin (BAF) treated recipient cells were measured and compared against the amount of miRNA found to present with Ago2 from siRILP treated cells. A relative decrease of Ago2 association of internalized miRNA was detected when cells were treated with Bafilomycin. Although total miRNA content release were comparable in both Bafilomycin and siRILP-treated cells against respective control groups. (H) Schematic illustration of the isolation procedure of endosomes from lysates of HeLa cells treated with Bafilomycin (*left panel*). Repeated ultracentrifugation were done to separate the endosomal compartment from released endosomal components in the reaction buffer after the desired reaction was done with endosomes isolated from Optiprep density gradient separated fractions positive for endosomal marker proteins. Effect of Proteinase K treatment on endosome associated Ago2. Western blot to show the sensitivity of Ago2 protein towards PK. Increasing concentration of Proteinase K was used alone or in presence of Triton X-100 (*right panel*). (I) Western blot analysis of the endosomal pellet and residual supernatant after separation of the endosomes isolated from fractions 2-4 of the 3-30% Optiprep^R^ gradient to confirm no ER (Calnexin) or Lysosomal (Lamp1) contamination in re-isolated endosomes used for pH change experiments described in panel J. (J-K) Schematic representation of the *in vitro* assay utilised to understand the role of pH change in release of internalized EV-entrapped miRNA. The endosomes were incubated with buffer of pH5 in presence and absence of FCCP, a protonophore, and then were separated from reaction buffer by ultracentrifugation (J, *left panel*). Quantification of miRNA-122 released after the incubation with FCCP in low pH buffer was estimated by qRT-PCR. Ct values of miR122 in the endosomal supernatant incubated with buffer of pH5 +/− FCCP (n=5) (J *right panel*). Time kinetics of release of miRNA into the supernatant from endosomes incubated with buffer of pH5+FCCP as determined by Real Time PCR (n=4) (K, *upper panel*).Pelleted membrane after the pH change reaction was performed was analyzed by western blot for endosmal protein Alix and HRS (K, *lower panel*).Western Blot of the supernatant and pellet of the FCCP treated endosomes was done to show the absence of endosomal membrane contamination in the supernatant recovered for miRNA estimation after treatment to account for the increase in miR-122 content in the supernatant. (L) Time dependent increase in miR-122 incorporation with endosome associated Ago2. Isolated endosomes, as described in panel J, was incubated with single stranded miRNA and were reisolated before Ago2 was extracted and amount of associated miRNAs was estimated. Levels of Ago2 in immunoprecipitated materials has been shown in Western blot. For statistical significance, minimum three independent experiments were considered in each case unless otherwise mentioned and error bars are represented as mean ± S.E.M. P-values were calculated by utilising Student’s t-test. ns: non-significant, *P < 0.05, **P < 0.01, ***P < 0.0001.

Bafilomycin is a proton pump blocker and it acts on endosomal maturation pathway by preventing the acidification of the endosomes prerequisite for its fusion with lysosome (Maroney et al, 2006). Bafilomycin’s effect has also been observed on the transfer of cargo from the early to the late endosomes (Bayer et al, 1998). Importantly, it has also been reported that low pH promotes the fusion of exosomal membrane with cell membranes (Parolini et al, 2009). Thus, it can be hypothesised that upon lowering the pH of endocytic vesicles, fusion between the exosomal membrane and internalized EV-membrane should occur. Since pH lowers as endosome matures, endosomal maturation should be an effective mechanism to trigger the release of EV-derived miRNAs in the proximity of endosomes and its loading on host Ago2 loading. To test this hypothesis, we blocked endosomal acidification with Bafilomycin. We detected an increase in EV-derived miRNA content upon treatment of recipient cells with Bafilomycin, concomitant to a decrease in the level of repression by internalized miRNAs (Fig 4C-F). This was consistent with the idea that Bafolimycin blocked the release of EV-derived miRNA from endosomes and would thus lead to the accumulation of EV-derived miRNA on endosomes. It was accompanied by a relative reduction in Ago2-bound internalized miR-122 levels compared to siRILP treated cells where RILP depletion also increases the total internalized miRNA content but with a concomitant increase in Ago2 association (Figure 4G).

To prove that low pH favors release of miRNA and miRNP formation with endosomal Ago2, we first proceed to check the localization of endosome associated Ago2 in mammalian cells. With isolated endosomal fractions from a Optiprep gradient separating the organelles, we found that the Ago2 present on endosome is sensitive to Proteinase K even in absence of any detergent like triton X-100. Therefore a large fraction of endosome localized Ago2 is present on the outer side of the endosomes and thus sensitive to Proteinase K (Figure 4H). Bafilomycin prevent the pH lowering of endosomes and thus endosomes isolated from Bafilomycin treated cells should have a lower capacity to release endocytosed EV-derived miRNAs, and this capacity should be restored by lowering endosomal pH. We purified endosomes after Bafilomycin treatment and incubated them in vitro in a buffer at pH5 with or without FCCP, a protonophore that can equalize the pH across biological membranes. We then followed the release of EV-derived miRNAs from endosomes after the *in vitro* reaction, by separating the endosome as pellet from the supernatant by ultracentrifugation. (Figure 4H, 4J). It was found that the exosomal miRNA content in the supernatant was increased in presence of FCCP and the increase has happened in a time dependent manner (Figure 4K). The endosomal pellets and supernatants were analysed by Western blot before and after the assay and this showed that a similar amount of endosomes was recovered with FCCP, ruling out that a change in endosomes themselves cause the changes (Figure 4I, K). These data thus have substantiated the idea of pH-dependent release of miRNA content from endosomes for a re-loading to membrane attached Ago2. This was furthur supported in an in vitro loading assay done with endosome associated Ago2. In this assay incubation of single-stranded miRNA with endosmes resulted in an time dependent miRNA-loading of Ago2-associated with endosomes (Figure 4L). These data thus have substantiated the idea of pH dependent release of miRNA content from endosomes for a re-loading to membrane attached Ago2. Since there are contact points between endosomes and the ER (Friedman et al, 2013) and endosomal Ago2 freshly loaded with EV-derived miRNAs could easily access the ER for its functioning.

### *Leishmania donovani (Ld)* prevents internalization of pro-inflammatory hepatic miR-122 in host macrophage

Next, we aimed at demonstrating the biological importance of endosomal uptake of EV-derived miRNA, and we looked for pathogen that would alter internalization of the EV-derived miRNA for its benefit. *Leishmania donovani* (*Ld*) is a biphasic pathogenic protozoan parasite that in its infective promastigote stage is transferred from the insect sand-fly reservoir to mammalian host and infects and resides inside tissue macrophages in the liver and spleen of the host mammals. *Ld* causes visceral leishmainasis (Murray et al, 2005) and is known to cause a robust change in endocytic pathway of the host cells and hijacks some components of the same pathway that are also found to be used for endocytic entry or exit of miRNAs (Chakrabarty & Bhattacharyya, 2017; Lievin-Le Moal & Loiseau, 2016). *Ld* is also known to affect miRNA machineries of host cells to ensure its proliferation (Ghosh et al, 2013). Going with the notion, close proximity of endosome with *Ld* containing parasitophorous vacuole and endosomes in infected macrophage cells were documented (Figure S4A-B). We explored the status of Rab proteins in *Ld* infected macrophage to document a decrease in endodsomal protein HRS and Rab7 in infected cells (Figure 5B and Figure S4C). miR-122 is known to have an pro-inflamatory role to play when transferred to macrophage cells (Ghosh et al, 2013). As observed, the over-expression of miR-122 caused an increase in the TNF-α level in RAW cells (Figure 5C) like what bacterial lipopolysaccaride (LPS) derived from gram negative bacterial membrane also did. Conversely, when *L.donovani* infection in RAW cells expressing miR-122 was followed, we found increased pro-inflammatory cytokine level and low anti-inflammatory cytokine levels upon miR-122 expression (Figure 5D, E). Interestingly, the infection level of respective cells was found to be reduced upon expression of miR-122 (Figure 5F, G). This suggests that the hepatic miR-122 has an immuno-protective role and when transferred via EVs released by hepatic cells could prevent infection of neighbouring liver macrophage cells that are the first target of invading pathogen in infection context (Ghosh et al, 2013; Momen-Heravi et al, 2015). In order to survive, *Ld* must have adopted strategies to prevent this EV-mediated miRNA transfer process to stop inflammatory response in the host cells upon interaction with miR-122 positive EVs. We hypothesized that *Ld* could hijack the inflammatory machinery of the host cell by preventing the miR-122 containing EV transfer to infected macrophages, and thus ensure survival of the pathogen. Consistent with the assumption, we found an increased production of pro-inflammatory cytokines and miR-155, the hallmark of inflammatory response in macrophage cells upon treatment with miR-122 positive EVs. But in cells infected with *Ld*, the uptake of miR-122 containing EVs was significantly compromised (Figure 5 H). As reported previously, *Ld* reduces the expression of HRS that is involved in endosome maturation (Figure 5B) however, we could not identify any major change in endosome number in *L.donovani* infected cells and neither the infection had any effect on Dynamin2 expression to account for the reduction in miR-122 entry (Figure S4A-C).

**Figure 5.**
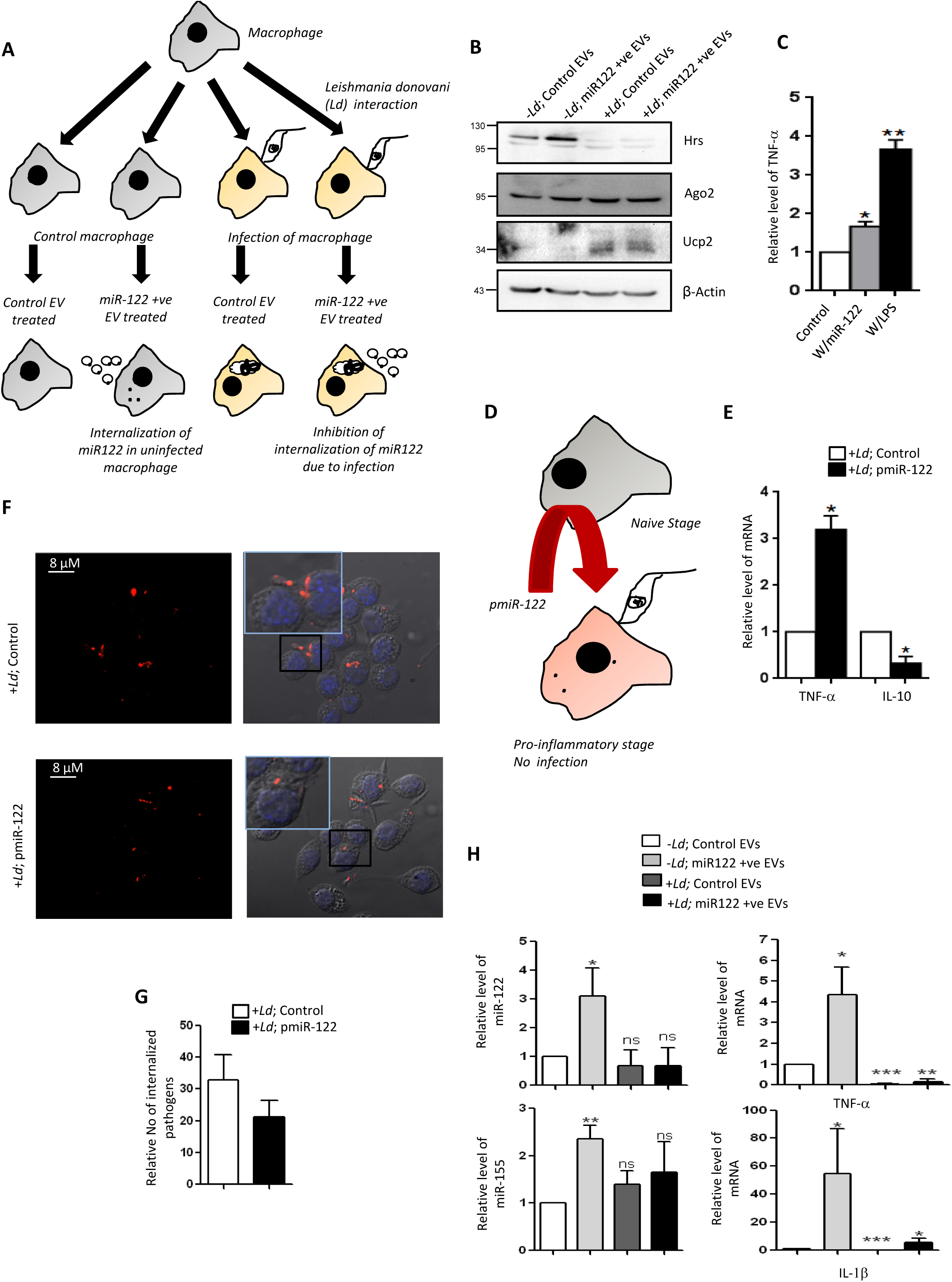
*Leishmania donovani* prevents internalization of EV-derived miR-122 in macrophage cells. (A) Schematic representation of experiments done with Leishmania infected or uninfected RAW264.7 cells treated with miR-122 containing EVs. Possible outcome of infection and miR-122 EV treatment has been hinted. (B) Western blot analysis of RAW264.7 cells to show the infection related upregulation of UCP2 and downregulation of HRS protein expression in RAW264.7 cells. (C) Level of TNF-α in RAW264.7 cells transfected with miR-122 expression plasmid to show the upregulation of proinflammatory cytokines in the cells on expression of miR-122. LPS was used as a positive control to induce TNF-α expression. (D) Scheme of miR-122 expression in RAW264.7 cells followed by infection with *Leishmania donovani* to detect its effect on proinflammatory cytokine production and on the level of infection in the 264.7 cells. (E) Relative levels of TNF-α and IL-10 in the miR-122 transfected RAW264.7 cells that are subsequently infected with *Ld* (n=3). (F, G) Level of infection in RAW264.7 cells transfected with miR-122 expressing pmiR122. Quantification of internalized pathogens (G) was done from microscopic images (n=100 cells, from four independent experiments) obtained for non miR-122 expressing and miR-122 expressing RAW264.7 cells (F). (H) Relative levels of internalized miR-122 in RAW246.7 cells which were either infected or uninfected with *Leishmania donovani* and treated or untreated with miR-122 containing EVs (*top left panel*). Relative level of endogenous miR-155 in RAW264.7 cells treated or untreated with miR-122 containing EVs and Leishmania is also shown (*bottom left panel*). Relative levels of pro-inflammatory cytokines TNF-alpha and IL-1β in response to miR-122 EV treatment or *Leishmania* infection and both (*right top and bottom panel*) (n=4). For statistical significance, minimum three independent experiments were considered unless otherwise mentioned and data are represented as mean ± S.E.M. P-values were calculated by utilising Student’s t-test. ns: non-significant, *P < 0.05, **P < 0.01, ***P < 0.0001.

### The mitochondrial protein Ucp2 is required for the internalization and activity of EV-derived miRNAs in recipient cells

*Ld* could target the mitochondrial dynamics and activity by inducing depolarization of mitochondria and reducing the mitochondria-ER-endosome interaction in infected cells (Chakrabarty & Bhattacharyya, 2017). We observed that mitochondrial uncoupler protein Ucp2 got upregulated upon *Ld* infection which might be the primary cause for the inhibition of the miRNA entry via EVs (Figure 5B). As mentioned, the mitochondrial depolarisation might play a role in affecting internalization of EV-associated miRNAs. The mitochondria can interact with several cellular compartments, like the ER and endosomes (Klecker et al, 2014; Todkar et al, 2019). In addition, the mitochondria can regulate miRNA activity by the presence of an active miRNA ribonucleoprotein complex (miRNP) present there (Wang & Springer, 2015). The Ucp2 is a mitochondrial uncoupling protein causing oxidative phosphorylation and ATP synthesis to uncouple by changing membrane potential across mitochondrial membranes. Hence, it was anticipated that uncoupling the mitochondrial potential can affect the internalization of EV-derived miRNAs in mammalian cells as it is known to affect endogenous miRNA activity (Chakrabarty and Bhattacharyya, 2017). FH-Ucp2 was expressed in the recipient HeLa cells and these cells were incubated with miR-122-containing exosomes. Ucp2 over-expression caused a disruption in the mitochondrial structure as it was observed microscopically (Figure 6A). The low level of internalization and consequently a lower repression activity of the transferred EV derived miRNA were observed in FH-Ucp2 expressing recipient HeLa cells (Figure 6B-D). It has been reported earlier that the loss of mitochondrial membrane potential is accompanied by reduced juxtaposition of endoplasmic reticulum (ER) and mitochondria (Chakrabarty & Bhattacharyya, 2017)(Figure S5A, S5B). Mfn2 is a protein responsible for mitochondrial tethering with ER. Mfn2 knockout mouse embryonic fibroblasts (MEFs) showed a decrease in EV associated miRNA internalization and Ago2 association compared to cells with Mfn2. Importantly, as reported earlier, the endogeneous miRNA miR-16 level was found to be increased in MFN2 depleted cells (Chakrabarty & Bhattacharyya, 2017) (Figure 6E-F) consistent with the previous finding on cellular miRNA content upon Ucp2 over-expression (Chakrabarty & Bhattacharyya, 2017).

**Figure 6.**
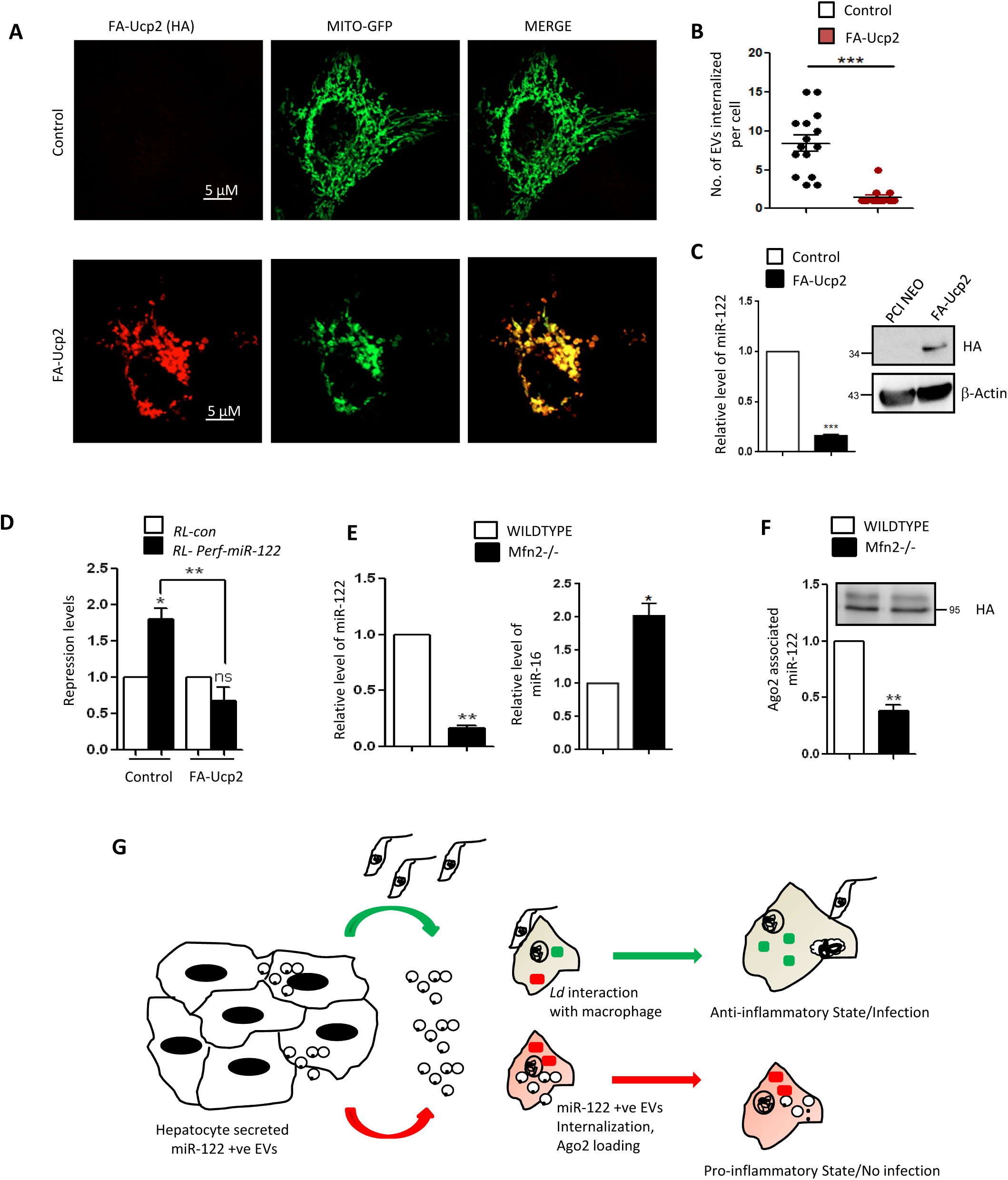
Mitochondrial activity is required for the internalization and repressive activity of EV-derived miRNAs in recipient cells. (A) Microscopic images to show the change in the structure of mitochondria on expression of mitochondrial uncoupler protein FH-Ucp2 in HeLa cells also expressing Mito-GFP. The FH-Ucp2 was detected by indirect immunoflurescence (red) and the mitochondria by GFP fluorescence (green). Scale bar 10 µm. (B) Microscopic analysis of internalization of CD63-GFP positive EVs in recipient HeLa cells transfected either with control or FH-Ucp2 expressing plasmids was done and graphical representation of quantification on EVs number in the recipient cells has been shown (n=15). (C) Relative level of internalized miR-122 via EV-mediated transfer in recipient HeLa cells transfected either with control or FH-Ucp2 expressing plasmids (n=3). (D) Repression levels of RL-perfmiR-122 in recipient cells transfected with control plasmid or FLAG-HA-Ucp2 and treated with miR-122 containing EVs. (E) Relative level of internalized miR-122 (*left panel*) and endogenous miR-16 levels (*right panel*) in Mfn2 wildtype and knockout mouse embryonic fibroblast (MEF) cells treated with EVs packed with miR-122. All quantification was done by qRT-PCR and U6 levels were used for normalization. (F) FH-Ago2 associated miR-122 levels in Mfn2 wild type and knockout FH-Ago2 expressing MEF cells co-cultured with miR-122 expressing HeLa cells. All quantification was done by qRT-PCR and Ago2 levels were used for normalization. (G) Pictorial representation of EV-mediated crosstalk between hepatocytes and macrophages. The miR-122 containing EVs released by hepatocytes can be transferred to macrophages that causes production of pro-inflammatory cytokines and could prevent *Ld* infection (red arrow). On the contrary, infected macrophages are unable to take up the miR-122 containing EVs due to upregulation of Ucp2 protein in cells infected with *Ld* and are tuned to have high production of anti-inflammatory cytokines (green arrow). For statistical significance, minimum three independent experiments were considered unless otherwise mentioned and data are represented as mean ± S.E.M. P-values were calculated by utilising Student’s t-test. ns: non-significant, *P < 0.05, **P < 0.01, ***P < 0.0001.

## Discussion

To interpret the interplay between *Leishmania* infection and EV-mediated miRNA entry pathway, we concluded a simplistic model of infection controlled EV-entry in macrophage cells (Figure 6G) where the hepatocyte released miR-122 containing EVs are to be internalised by surrounding macrophages. The recipient cells’ endosomal pathway was used for effective entry of the EV-derived miRNAs. Hepatocyte specific miR-122 ensures activation of recipient macrophage and pro-inflammatory cytokine production in liver resident macrophages to prevent tissue infection. However, in the infected macrophages, the pathogen prevents the internalization of EVs by both disrupting HRS and the mitochondrial membrane potential. This prevents uptake of EV-derived miRNAs and leads to an upregulation of the anti-inflammatory response that facilitates sustained infection.

Extracellular vesicles or EVs have recently been established as another means of intercellular communication apart from that mediated through cell junctions and secretary molecules. Earlier EVs were collectively considered as vehicles of waste disposal and to carry cellular junk out of the cells. There have been ample evidences in support of role of EVs in carrying genetic materials and other biologically important cargos to other target cells to elicit a response there (Au Yeung et al, 2016; Cossetti et al, 2014; Thery et al, 2002; Valadi et al, 2007). Present concept, strongly supported by experimental data, suggests that the miRNA mediated gene silencing can be achieved via extracellular vesicle induced crosstalk between cells (Chiba et al, 2012; Zhang et al, 2015). Recent studies have also shown the existence of a difference in the expression profiles of circulating miRNAs in the blood or other body fluids of healthy and diseased patients which can serve as the potential biomarkers for disease diagnosis. However a major limitation in EV-miRNA field is the deficit in the understanding of the mechanistic aspect of functional internalization of the EV-carried miRNAs into recipient cells. Our findings shed light on the mechanism of transfer of microRNA between mammalian cells.

Here in this study we have elucidated how the functional miRNA can be transferred from donor to recipient cells via EVs. The transferred miRNA does not affect the pre-exisiting endogenous miRNA levels as well as their activities. We have observed that the exosomal miRNA is transferred primarily in a Dicer independent manner in its mature single stranded form. This finding is consistent with the fact that the precursor miRNAs get processed to mature miRNAs due to the presence of RISC Loading Complex in exosomes (Melo et al, 2014). Previous studies on stabilization of the circulating miRNAs by the Ago2 protein has been reported earlier (Turchinovich et al, 2011). Also EV-sorting of Ago2 has been reported to stabilize miRNAs in exosomes or microvesicles (Beltrami et al, 2015; Lv et al, 2014; McKenzie et al, 2016). Despite the earlier claim on presence of Ago2 in the EVs, recent publication question the presence of protein components of miRNP machinery in mammalian cell derived EVs (Jeppesen et al, 2019). Therefore, it is most likely that single stranded miRNAs may travel from one cell to other via EVs in a Ago2 unbound form. We have observed that the EV-derived miRNA get bound to the recipient cell Ago2 that drives the sustained function of the transferred miRNA in mammalian cells. However, a possible role of donor cell Ago2 in this process can’t be ruled out.

Compartmentalization of miRNA function and mRNA degradation is a much studied aspect of miRNA biology. Recent findings suggest the involvement of different sub-cellular organelles like the Endoplasmic Reticulum and the endosomes in miRNA mediated mRNA repression (Bose et al, 2017; Lee et al, 2009; Li et al, 2013; Siomi & Siomi, 2009). Previous evidences establish rER as the site of assembly of the mRNA silencing machinery (Stalder et al, 2013). It is also the site of mRNA-miRNP interaction (Barman & Bhattacharyya, 2015). These repressed mRNAs have been found to be relocated subsequently to the multivesicular bodies (MVBs) where they get released from Ago2 and subsequently get degraded (Bose & Bhattacharyya, 2016). Internalization of the EV-derived miRNA is also found to be compartmentalized to enable its effective functioning. We have demonstrated that the uptake of the exosomal miRNA is dependent on Dynamin2, a GTP binding protein involved in the endocytosis pathway. It utilises the endosomal pathway to reach the Endoplasmic Reticulum of the recipient cell. The compartmentalization of the exosomal miRNA to ER suggests adaptability of the same to elicit a repressive activity in the recipient cell. We have found that the knockdown of the endosomal pathway effector protein RILP leads to accumulation of the internalized miRNA in the recipient cell that cannot repress its target genes. Subcellular localization assays revealed a differential localization and Ago2 association of the transferred miRNAs on knockdown of RILP. These results explain why the internalized miRNA cannot be functional as it fails to reach the ER for its activity in endosome maturation defective cells.

The recent advancement in 3D electron microscopy has helped us to reveal ER and endosome binding and it is known that when the endosomes transit through steps of maturation, they become more tightly associated with the ER (Friedman et al, 2013). Thus, knockdown of the endosomal pathway proteins prevent association of the internalized miRNA with the ER. The pH of the endosomal vesicles plays a role in release of cargo molecules to the late endosomal and lysosome attached cytoplasmic components. It has been observed that endosomal pH and hence maturation plays a role in uncoating of viral particles for their infectivity (Li et al, 2014). Also low pH can influence the release and uptake of exosomes by fusion in cancer cells (Parolini et al, 2009). Hence, an *in vitro* assay was done to validate that the change in pH during the endosomal maturation may play a role in release of the late endosomal contents into the ER via the contact points. Loading of single stranded miRNA to Ago2 attached to endosome membranes strongly support the concept of endosomal miRNP formation with memebrane bound Ago2 and internalized miRNAs.

*Leishmania donovani (Ld)*, the causative agent of visceral leishmaniasis, is known to upregulate anti-inflammatory cytokines in infected macrophages to establish infection. Our observations elucidate the mechanism how the kinetoplastid parasite can restrict the entry of miR-122 containing EVs in infected macrophages by mitochondrial depolarisation. miRNA-122 can activate the pro-inflammatory cytokines in the macrophages and we have documented how miR-122 expressing RAW cells can curtail infection by upregulation of pro-inflammatory cytokine, TNF-α and downregulation of anti-inflammatory cytokine, IL-10. These have been found to be consistent with the observation in visceral leishmaniasis. In *Ld* infected mouse liver, restoration of miR-122 levels can reduce parasite burden and increase the serum cholesterol levels (Ghosh et al, 2013). In order to revalidate the mitochondrial depolarisation in *Ld* infected cells as the cause of inhibition of EV-mediated miRNA entry in RAW cells, we expressed Ucp2 in recipient cells to document a decrease in uptake and activity of the EV-derived miRNAs. Mitochondrial disruption may cause a break in the contact of mitochondria with the ER hence hampering the transfer of the miRNA to the ER or driving it away for lysosomal degradation or out of the cell by EVs. This work provides an insight into the machineries that the EV-derived miRNAs use for entering the recipient cell. However, with depletion of Ucp2 by siRNAs in receipinet cells infected with *Ld* we could not rescue the internalization of miR-122 containing EVs. (data not shown). This may be explained easily considering the global effect *Ld* infection has on endocytic pathway by lowering HRS or Rab7 protein expression detected in infected cells. Thus *Leishmania* adopts multiple paths apart from mitochondrial depolization to achieve the inactivation and endocytic entry of pro-inflammatory miR-122 in infected macrophages. In this context we have reported, in our recent work, how the Leismania by restricting the expression of miRNA exporter protein HuR can delimit the miRNA export and macrophage activation process (Goswami et al, 2020). Therefore targeting endosomal miRNA entry and its re-activation in pathogen internalized cell is a interesting additional machinery for the pathogen to ensure robust anti-inflammatory response in infection context.

The miRNA uptake via EVs ensures sensing of extra-cellular status and its coupling with gene repression machineries of respective cells. By responding to the cellular miRNA levels, the cellular miRNA uptake machineries may balance the cellular miRNA content to balance the gene repression. Therefore this may be considered as a salvage pathway for *de novo* miRNA formation in mammalian cells to buffer the cellular miRNA content.

## MATERIALS AND METHODS

### Cell culture and parasite infection

Human HeLa, Huh7 as well as mouse embryonic fibroblast cells (WT/ MFN2-/-) were cultured in Dulbecco’s Modified Eagle’s medium (DMEM; Gibco-BRL) supplemented with 2mM L-glutamine and 10% heat-inactivated fetal bovine serum (FBS). Exosome depleted growth media was made with DMEM and 10% exosome depleted FBS. For depletion of exosomes from FBS, the FBS was ultra-centrifuged at 1,00,000X g for 4 hours and then added to the DMEM. Characterisation of EV depletion was done as reported previously (Mukherjee et al, 2016). Plasmid transfections were done with Lipofectamine 2000 (Invitrogen) following manufacturer’s instruction. All plasmid constructs used were reported previously (Ghosh et al, 2015)

The murine macrophage RAW 264.7 cells were cultured in RPMI 1640 medium (Gibco) supplemented with 2mM L-glutamine, 0.5% β-mercaptoethanol and 10% heat-inactivated fetal bovine serum.

For parasitic infections of RAW 264.7 cells, 2nd - 4th passage *in vitro* cultures of *Leishmania donovani* AG83 promastigotes, grown in M199 medium supplemented with 10% FCS, were used to infect cells with a 10:1 ratio for all experiments for 6 hours or 16 hours as mentioned for specific experiments.

### siRNA and plasmid transfections

siRNAs against different endocytic pathway proteins (Rab 5, Rab 7 and RILP) and siDicer and siAgo2 were purchased from Dharmacon. siRNA transfection was done using RNAi Max (Invitrogen) according to manufacturer’s instructions. HeLa cells were transfected with 15 pmoles of siRNA per well of a 24 well plate. The siRNA transfected cells were split 48-72 hours post transfection for proper knockdown. List of different siRNAs used are mentioned in supplementary table S2.

For exosome isolation, donor HeLa cells were transfected with 1 μg pmiR122 per well of a 6 well plate. pmiR122 plasmid, a plasmid encoding the precursor miR122, was described elsewhere (Ghosh et al, 2013)

### Exosomes isolation and characterisation

HeLa cells transfected with pmiR122 were split 24 hours post transfection to 90 mm plates and incubated for 24hours at 70-80% confluency. These cell culture supernatants were subjected to EV isolation as described previously with minor modifications (Thery et al, 2006) For EV isolation, cells were grown in exosome depleted media to prevent any background from exosomes or EVs present normally in FBS. Briefly, the cell supernatant (exosome depleted growth media which was used to culture the donor miR-122 expressing HeLa cells) was centrifuged at 2,000X g for 15 minutes to remove cell debris. Next the cell supernatant was collected and centrifuged at 10,000X g for 30 minutes. The supernatant that was obtained was passed through 0.22μm filter unit to further clear it. This was followed by ultracentrifugation of the supernatant at 1,00,000X g for 90 minutes. After ultracentrifugation, the pellet was resuspended in exosome depleted growth media and added to recipient cells.

For characterisation of EVs by Atomic Force Microscopy, the EVs were isolated on a 30% sucrose cushion and the layer of exosomes were further washed and pelleted with 1X PBS by ultra-centrifugation at 1,00,000X g for 90 minutes and resuspended in 1ml PBS. Then 5ul of the EV suspension was placed onto mica sheet and dried for 15 minutes. The sample slides were then gently washed with autoclaved MilliQ water to remove molecules that were not firmly attached to the mica and dried again. AAC mode AFM was performed using a Pico plus 5500 ILM AFM (Agilent Technologies USA) with a piezoscanner maximum range of 9 µm. Micro fabricated silicon cantilevers of 225 μm in length were used from Nano sensors, USA. Images were processed by flatten using Pico view1.1 version software (Agilent Technologies, USA).

For Nanoparticle tracking analysis (NTA), EVs were resuspended in 1ml PBS. EVs were 10-fold diluted and 1ml of diluted EVs were injected in to the sample chamber of Nanoparticle tracker (Nanosight NS300).

### Contact co-culture model

Donor HeLa cells were transfected with pmiR122 while recipient cells were transfected either with RLcon, RL-perfmiR-122 or FLAG-HA-Ago2 plasmids depending on whether luciferase assay or immunoprecipitation (IP) assays were done. After transfection, the donor and recipient cells were seeded in equal amounts and allowed to interact for 24 hours after which either luciferase assay or IP was done. IP was followed by quantification of the amount of miRNA transferred to the recipient Ago2 in the contact co-culture system while luciferase assay depicted the amount of RL reporter repression in recipient cells in presence of internalized miRNA from donor cells.

### Luciferase assay

The plasmids RL control and RL-perfect-miR122 substrates were procured as a kind gift from Dr. Witold Fillipowicz. For miRNA repression assays, 30 ng of Renilla luciferase (RL) reporter plasmids (both RL con and RL perfect 122) with 300 ng of firefly luciferase (FL) plasmid were co-transfected per well of a 12-well plate. RL and FL activities were measured using a Dual-Luciferase Assay Kit (Promega, Madison, WI) following the supplier’s protocol on a VICTOR X3 Plate Reader (PerkinElmer, Waltham, MA). The RL expression levels for reporter and control were normalized against Firefly levels. These normalized values were then used to calculate fold repression as the ratio of normalized control to reporter RL values, i.e, RLcon/RLperfect.

### Immunoprecipitation assay

For immunoprecipitation of Ago2, HeLa cells were transfected with FLAG-HA tagged Ago2 plasmid. Briefly, Protein G agarose beads (Invitrogen) or FLAG-M2 agarose beads (Sigma) was used for Flag tagged Ago2 IP. For HA, beads were blocked with 5% BSA in lysis buffer for 1hr followed by antibody incubation (1:50 dilution) for 4 hours at 4°C. For IP reactions, HeLa cells were lysed in Lysis buffer (20 mM Tris-HCl, pH 7.5, 150 mM KCl, 5 mM MgCl2, 1 mM DTT) with 0.5% Triton X-100, 0.5% sodium deoxycholate and 1X EDTA-free protease inhibitor cocktail (Roche) for 20 min at 4°C. The lysates, clarified by centrifugation at 3,000X g for 10 minutes, were incubated with HA antibody pre-bound Protein G Agarose bead or pre-blocked anti-FLAG M2 beads and rotated overnight at 4°C. Subsequently, the beads were washed thrice with 1X IP buffer (20 mM Tris-HCl pH 7.5,150 mM KCl, 5 mM MgCl2, 1 mM DTT), and separated into two halves for RNA and protein analysis from the bound Ago2 on the beads.

### Optiprep density gradient ultra-centrifugation

Optiprep™ (Sigma-Aldrich, USA) was used to prepare a 3-30%continuous gradient in a buffer containing 78 mM KCl, 4 mM MgCl_2_, 8.4 mM CaCl_2_, 10 mM EGTA, 50 mM Hepes (pH 7.0) for separation of subcellular organelles. HeLa cells were washed with PBS and homogenized with a Dounce homogenizer in a buffer containing 0.25 M sucrose,78 mM KCl,4 mM MgCl_2_,8.4 mM CaCl_2_,10 mM EGTA, 50 mM Hepes pH 7.0 in addition to 100 μg/ml of Cycloheximide, 5 mM Vanadyl Ribonucleoside Complex (VRC) (Sigma Aldrich), 0.5 mM DTT and 1X Protease Inhibitor. The lysate was clarified by centrifugation at 1,000X g for 5 minutes two times and layered on top of the prepared gradient. The tubes were centrifuged at 1,33,000X g for 5 hrs for separation of gradient and ten fractions were collected by aspiration from the top. The fractions were pooled accordingly for subsequent analysis of proteins and RNA. For immunoprecipitation of FLAG-HA-Ago2 from the fractions, the pooled fractions were lysed with lysis buffer for 20 minutes at 4°C. The lysate was clarified by centrifugation at 16,000X g for 5 minutes and incubated with FLAG beads overnight followed by RNA and protein analysis of the Ago2 associated with the beads.

### Microscopic analysis of EV entry into recipient cells

To detect the uptake of EVs by microscopy, the donor HeLa cells were transfected with CD63-GFP expression plasmid encoding a GFP tagged exosomal tetraspanin. The supernatant of these cells was used for isolation of EVs by Exosome Isolation Reagent (Thermo Scientific) from cell culture medium. The supernatant was cleared of cell debris at 2,000X g for 30 mins followed by addition of this reagent which is incubated overnight at 4°C. On the following day the sup is centrifuged at 10,000X g for 60 mins to pellet the EVs. These are added to the recipient cells seeded on coverslips at different conditions to observe under the microscope. Cells were fixed using 4% para-formaldehyde for 20mins. For detection proteins like FLAG-HA-UCP2 or β-tubulin, blocking and permeabilization was done by 1% BSA, 10% goat serum and 0.1% triton X-100 for 30 mins. The Anti-HA (Rat) and Anti-β-tubulin (Mouse) were used at 1:1000 dilution. Secondary Alexafluor anti-rat and anti-mouse were 568 dye fluorochrome tagged and were used at 1:500 dilutions. To detect the Leishmanial infection in RAW cells, anti-GP63 (Mouse) was used in 1:1000 dilution followed by anti-mouse 568 dye at 1:500 dilution. For calculating % of infected cells, minimum of 100 cells were counted. All the microscopic detection was done using Zeiss LSM800 followed by analysis using Imaris software.

### *In vitro* miRNA release assay

To see the effect of pH on release of miRNA from endocytic vesicles, an *in vitro* assay was done. For this the recipient cells were incubated with 25 nM bafilomycin for 17 hours and treated with miR-122 positive EVs. The cell lysates were subjected to Optiprep density gradient ultracentrifugation the early endosomal marker positive fractions 2, 3, 4 were collected and diluted with buffer. This was followed by further purification of the endosome by ultracentrifugatiom at 1,33,000X g for 2.5 hours. The endosome pellet was then resuspended in 100 µl Tris-HCl buffer of pH5 with or without FCCP. The endosomal buffer was then incubated for 30 minutes at 37°C. The reaction was stopped by transferring the mixture at 4°C. This was further diluted with a buffer containing 78 mM KCl, 4 mM MgCl_2_, 8.4 mM CaCl_2_, 10 mM EGTA, 50 mM Hepes (pH 7.0). Then again, the mixture was ultracentrifuged at 1,33,000X g for 2.5 hours to pellet the endosome and then the supernatant was used to isolate RNA to see the amount of miRNA released from the vesicles. The remaining supernatant and pellet were used for Western Blot analysis of marker proteins to rule out the contamination of endosomes in the supernatant.

### Proteinase K protection assay

In order to detect whether the Ago2 protein is present on the outer or inner surface of the endosomal membranes, a ProteinaseK assay was done. Briefly, the HeLa cells were lysed and loaded on Optiprep density gradient and ultra-centrifuged as mentioned above. Then the endosome enriched fractions 2, 3, 4 were collected. The endosomal fractions were then incubated with 0, 5, 10, 20 ng/µl Proteinase K without TritonX and 20 ng/µl Proteinase K with TritonX for 30 minutes at 37 °C. After the reaction, the solutions were subjected to methanol chloroform protein precipitation. Then Western Blotting for Ago2 was done to see whether the Ago2 is present on the outer surface and Proteinase K is sufficient enough to disrupt the protein or is the Ago2 localized inside the endosomal compartment and Triton X-100 is needed to disrupt the membrane to enable the action of the ProteinaseK on Ago2.

### Polysome isolation

For total polysome isolation, HeLa cells were lysed in a buffer containing 10 mM HEPES pH 8.0, 25 mM KCl, 5 mM MgCl2, 1 mM DTT, 5 mM VRC, 1% Triton X-100, 1% sodium deoxycholate and 1 × EDTA-free protease inhibitor cocktail (Roche) supplemented with Cycloheximide (100 μg/ml; Calbiochem). The lysate was cleared at 3,000X g for 10 min followed by another round of pre-clearing at 20,000X g for 10 min at 4 °C. The clarified lysate was loaded on a 30% sucrose cushion and ultracentrifuged at 100,000X g for 1 h at 4 °C. The sucrose cushion was washed with a buffer (10 mM HEPES pH 8.0, 25 mM KCl, 5 mM MgCl2, 1 mM DTT), ultracentrifuged for additional 30 min and the polysomal pellet was finally resuspended in polysome buffer (10 mM HEPES pH 8.0, 25 mM KCl, 5 mM MgCl2, 1 mM DTT, 5 mM VRC, 1 × EDTA-free protease inhibitor cocktail) for protein and RNA isolation and estimation.

### Immunoblotting

The samples (cell lysates, membrane fractions, immunoprecipitated proteins) were subjected to SDS-polyacrylamide gel electrophoresis,transferred to PVDF nylon membranes and probed with specific antibodies for a minimum 16 hrs at 4°C. Following overnight incubation with antibody, membranes were washed and incubated at room temperature for 1 hr with secondary antibodies conjugated with horseradish peroxidase (1:8000 dilutions).Imaging of all western blots was performed using an UVP BioImager 600system equipped with VisionWorks Life Science software (UVP) V6.80. Antibody dilutions for HA, Rab5, Rab7, Calnexin, Lamp1 were 1:1000 while dilution for GAPDH was used at 1:50000.

### RNA Isolation and Real Time PCR

Total RNA is isolated by using TriZol or TriZol LS reagent (Invitrogen) according to the manufacturer’s protocol. MiRNA assays by real time PCR was performed using specific primers for human let-7a (assay ID 000377), human miR-122 (assay ID 000445), human miR-122* (assay ID 002130), human miR-16 (assay ID 000391). U6 snRNA (assay ID 001973) was used as an endogenous control. Real time analyses by two-step RT-PCR was performed for quantification of miRNA levels on Bio-Rad CFX96TM real time system using Applied Biosystems Taqman chemistry-based miRNA assay system. Cycles were set according to manufacter’s protocol. Samples were analyzed in triplicates. The comparative Ct method which included normalization by the U6 snRNA was used for relative quantification.

### *In vitro* endosomal Ago2 miRNA loading assay

To analyse the amount of miRNA bound to Ago2 in endocytic vesicles, an *in vitro* assay was done. For this FLAG-HA-HEK293 cell lysate was subjected to Optiprep density gradient ultracentrifugation as mentioned earlier and the early endosomal fractions 2, 3, 4 were collected and diluted with buffer 78 mM KCl, 4 mM MgCl_2_, 8.4 mM CaCl_2_, 10 mM EGTA, 50 mM Hepes (pH 7.0). This was followed by further purification of the endosome by ultracentrifugation for 2.5 hours. The endosome pellet was then resuspended in 50 µl buffer containing 0.25 M sucrose,78 mM KCl,4 mM MgCl_2_,8.4 mM CaCl_2_,10 mM EGTA, 50 mM Hepes pH 7.0, 100 μg/ml of Cycloheximide, 0.5 mM DTT, RNase Inhibitor (Applied Biosystem) and 1X Protease Inhibitor. These endosomal suspensions were incubated with 500 fmoles miR-122 for either 0, 15 or 30 minutes at 37°C. The reaction was stopped by further dilution with the same endosome resuspension buffer. The endosomes were lysed in a Lysis buffer comprising of 20 mM Tris-HCl, pH 7.5, 150 mM KCl, 5 mM MgCl2, 1 mM DTT with 1% Triton X-100 and 1X Protease Inhibitor for 20 min at 4°C. The endosomes were then sonicated with three pulses for 10 seconds each followed by centrifugation at 16,000X g for 5 minutes at 4°C to clear the lysate. This was then loaded onto FLAG beads for overnight immunoprecipitation of the cellular Ago2. The beads were washed on the following day and the beads were divided for analysis of protein and Ago2 associated RNA. Amount of miRNA was normalized against amount of Ago2 isolated.

### Statistical Analysis

All graphs and statistical significance was calculated using GraphPad Prism 5.00 (GraphPad, San Diego). Experiments were done for minimum of three times unless otherwise mentioned. Non-parametric Student’s t-test was used for determination of P-values.

## Authors Contributions

BG has done all the experiments. SNB conceptualize the idea, planned the experiments, analyzed the data and wrote the manuscript with BG. EB has helped in planning microscopy experiments.

## Acknowledgements

We acknowledge Witold Filipowicz for different plasmids constructs. SNB is supported by The Swarnajayanti Fellowship (DST/SJF/LSA-03/2014-15) from Dept. of Science and Technology, Govt. of India, while BG acknowledges support from Council of Scientific and Industrial Research, Govt. of India for Fellowship and CEFIPRA for Raman-Charpak travel exchange fellowship. The work was supported by funds from High Risk High Reward Project Grant, Dept. of Science and Technology, Govt. of India. The work also received support from a High Risk High Reward Grant (HRR/2016/000093) from Dept. of Science and Technology, Govt. of India.

## Conflict of interest statement

The authors declare no conflict of interest

## Supplemental Figure Legends

**Supplemental Figure S1.**
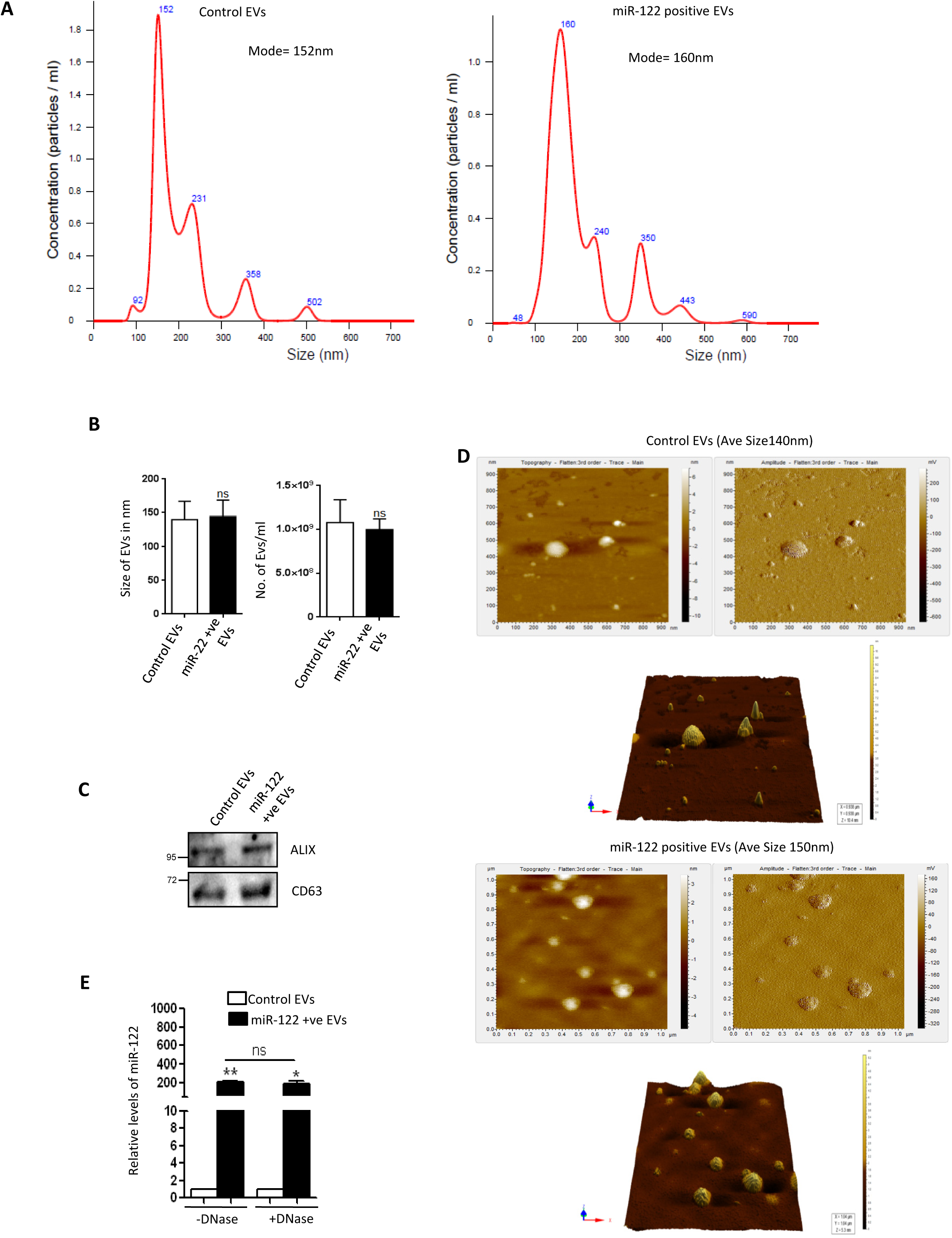
Characterisation of EVs released by miR-122 expressing donor HeLa cells. (A) Nanoparticle tracking analysis (NTA) to show similarity in the size of EVs released by untransfected and miR-122 expression plasmid transfected HeLa cells. Following EV isolation by ultracentrifugation of pre-cleared cell culture supernatants on sucrose cushion at 1,00,000X g, they were subjected to NTA analysis (B) Graphical representation of the size (*left panel*) and number of EVs released (*right panel*) as observed in NTA between untransfected control and miR-122 transfected HeLa cells. (C) Western blot data to show the protein levels of the EV marker proteins tetraspanin CD63 and Alix in EVs isolated from cell supernatants of control and miR-122 expressing HeLa cells. (D) Tapping mode Atomic Force Microscopic (AFM) images of EVs from control and miR-122 expressing cells showing round morphology of EVs. Middle panels shows 3-D image of the EVs. (E) Effect of DNase on miR-122 content with EVs. The isolated RNA from control and miR-122 containing EVs were subjected to RNase Free DNase I treatment followed by qRT-PCR to estimate the amount of miR-122 detected with or without DNaseI treatment. For statistical significance, minimum three independent experiments were considered unless otherwise mentioned and error bars are represented as ± S.E.M. P-values were calculated by utilising Student’s t-test. ns: non-significant, *P < 0.05, **P < 0.01, ***P < 0.0001.

**Supplemental Figure S2.**
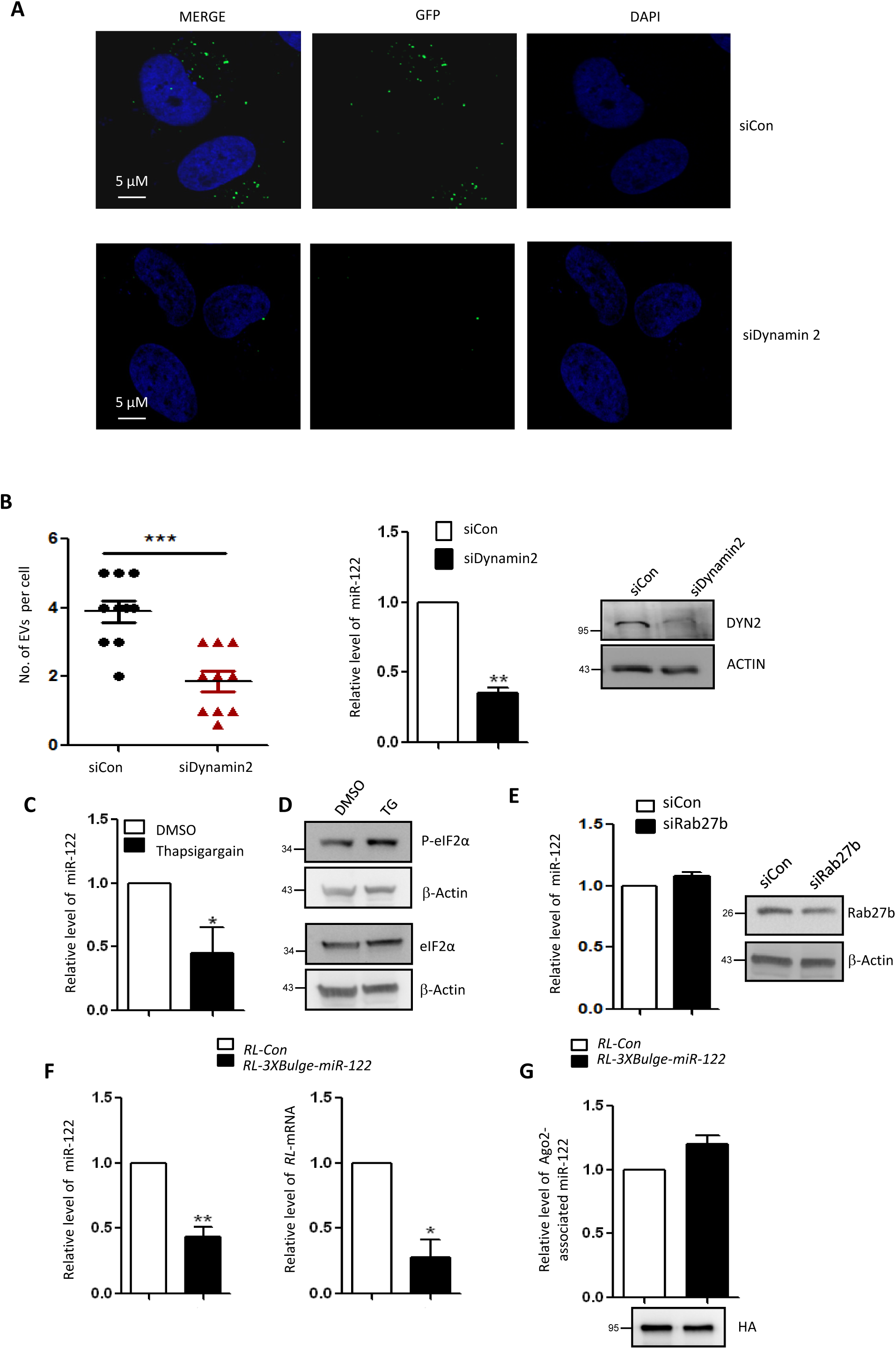
Effect of Dynamin2 depletion and target RNA availability on internalization of miR-122 via EVs. (A) Microscopic analysis of internalization of CD63-GFP positive EVs in recipient cells transfected with siCON and siDynamin2. (B) Quantification of internalization of CD63-GFP containing EVs in recipient cells transfected with siCON and siDynamin2 (*left panel, n=10*). Relative levels of internalized miR-122 in cells transfected with siCON and siDynamin2 by Real Time PCR (*middle panel*). U6 levels were used for normalization. Western blot analysis was used to check the knockdown of Dynamin2 in recipient HeLa cells (*right panel*). (C) Relative levels of internalized miR-122 in recipient cells treated with miR-122 positive EVs and exposed either to DMSO or Thapsigargin, an ER stress inducer. (D) Western blots of phospho-eIF2α and eIF2α in recipient cells treated with DMSO or Thapsigargin to show induction of stress in Thapsigargin treated cells. (E) Amount of miR-122 transferred via EVs to recipient cells transfected with siCon or siRab27b. Rab27a is a protein which is required for the secretion of miRNAs via EVs (Ostrowski et al., 2010). (F) Relative levels of internalized miR-122 (*left panel*) and substrate RL mRNA levels (*right panel*) in recipient HeLa cells transfected with RL-con or RL-3XBulge-miR122 after treatment for 16h with miR-122 positive EVs. The effect of presence of substrate on transferred RNA content and activity were measured. (G) FH-Ago2 associated miR-122 in recipient cells expressing either RL-con or RL-3XBulge-miR-122 mRNAs along with FH-Ago2. Recipient cells were co-cultured with miR-122 expressing donor HeLa cells. Amount of miR-122 bound to FH-Ago2 was measured in the presence of a substrate RL-3XBulge-miR-122 mRNA or control RL mRNA (n=5). For statistical significance, minimum three independent experiments were considered unless otherwise mentioned and error bars are represented as ± S.E.M. P-values were calculated by utilising Student’s t-test. ns: non-significant, *P < 0.05, **P < 0.01, ***P < 0.0001.

**Supplemental Figure S3.**
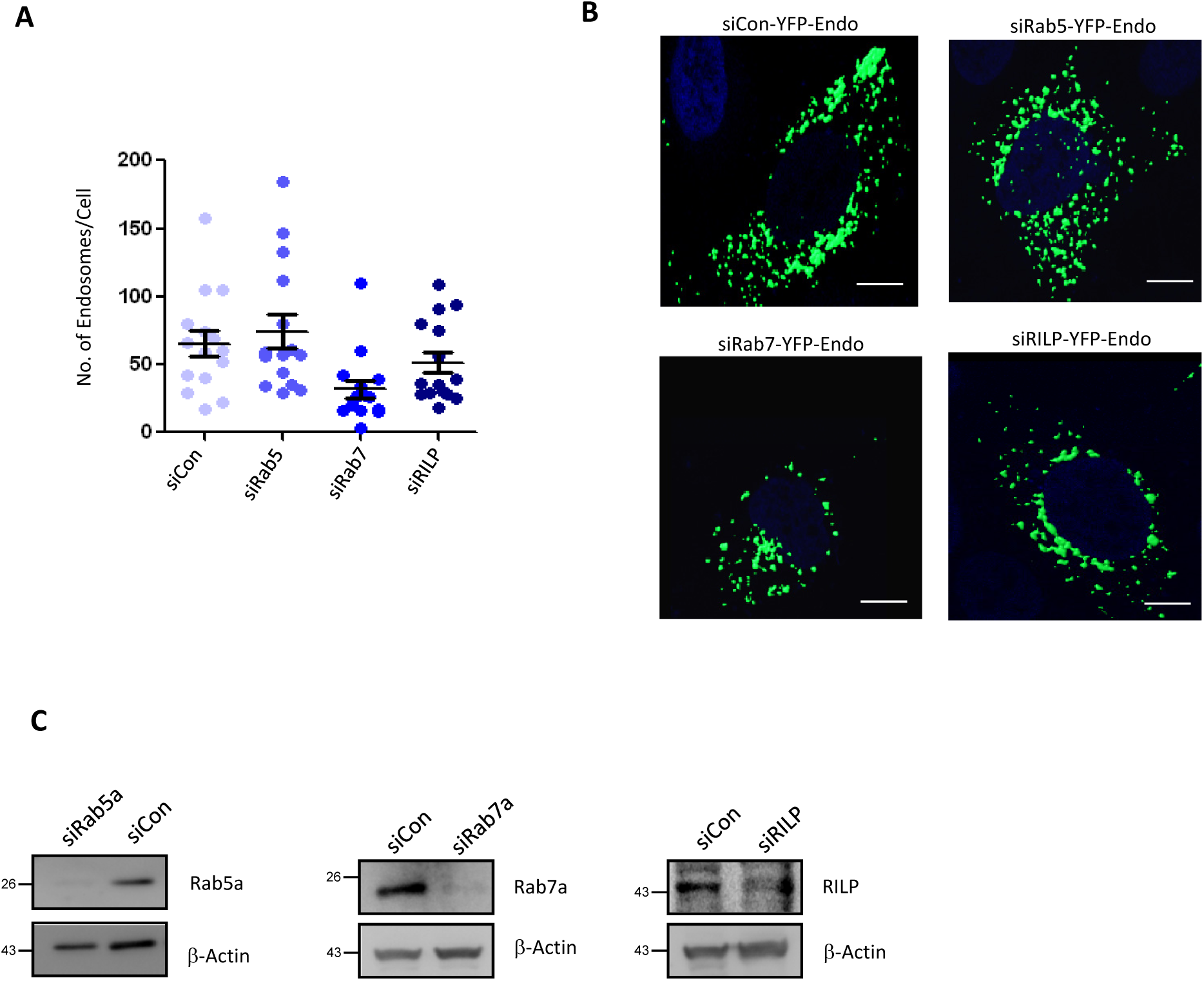
Effect of knockdown of endocytic pathway proteins on cellular endosomal compartment. (A) Estimation of number of endosomes in cell transfected with siCON, siRab5, siRab7, siRILP. The cells were co-transfected with YFP-Endo to label the endosomes. The number of endosomes per cell was counted using the Imaris software (n=15). (B) Representative images of the HeLa cells co-transfected with YFP-Endo and either siCON, siRab5, siRab7, siRILP. The endosomes are denoted in green (YFP-Endo). DAPI (blue) was used to stain the nucleus. Scale bar 10 µm. (C) Western blot analysis to check the knockdown of Rab proteins (Rab5, Rab7, and RILP) by siRNA transfection.

**Supplemental Figure S4.**
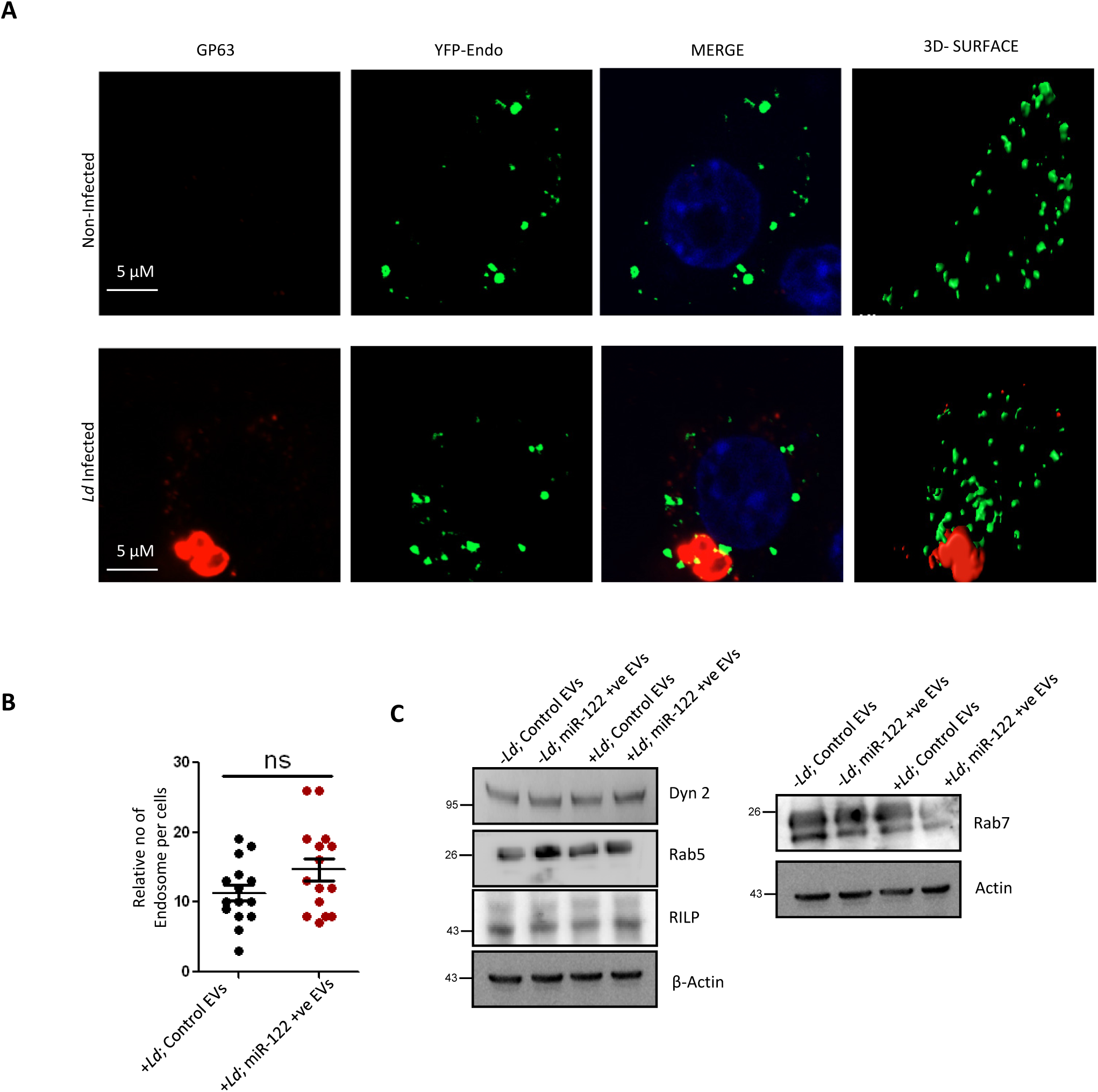
Effect of Leishmania infection on endosomes of RAW264.7 cells. (A) RAW cells, either *Leishmania* infected or naive, were imaged for the endosomes (marked with YFP-Endo, green) and Leishmanial GP63 protein (marked with red). (B) Quantification of the number of endosomes per cell as observed in both uninfected and infected RAW264.7 cells (n=15). (C) Western blots to show the levels of different endocytic pathway component proteins (Dynamin2, Rab5, Rab7, and RILP) under different conditions of infection and EV-treatment in RAW264.7 cells.

**Supplemental Figure S5.**
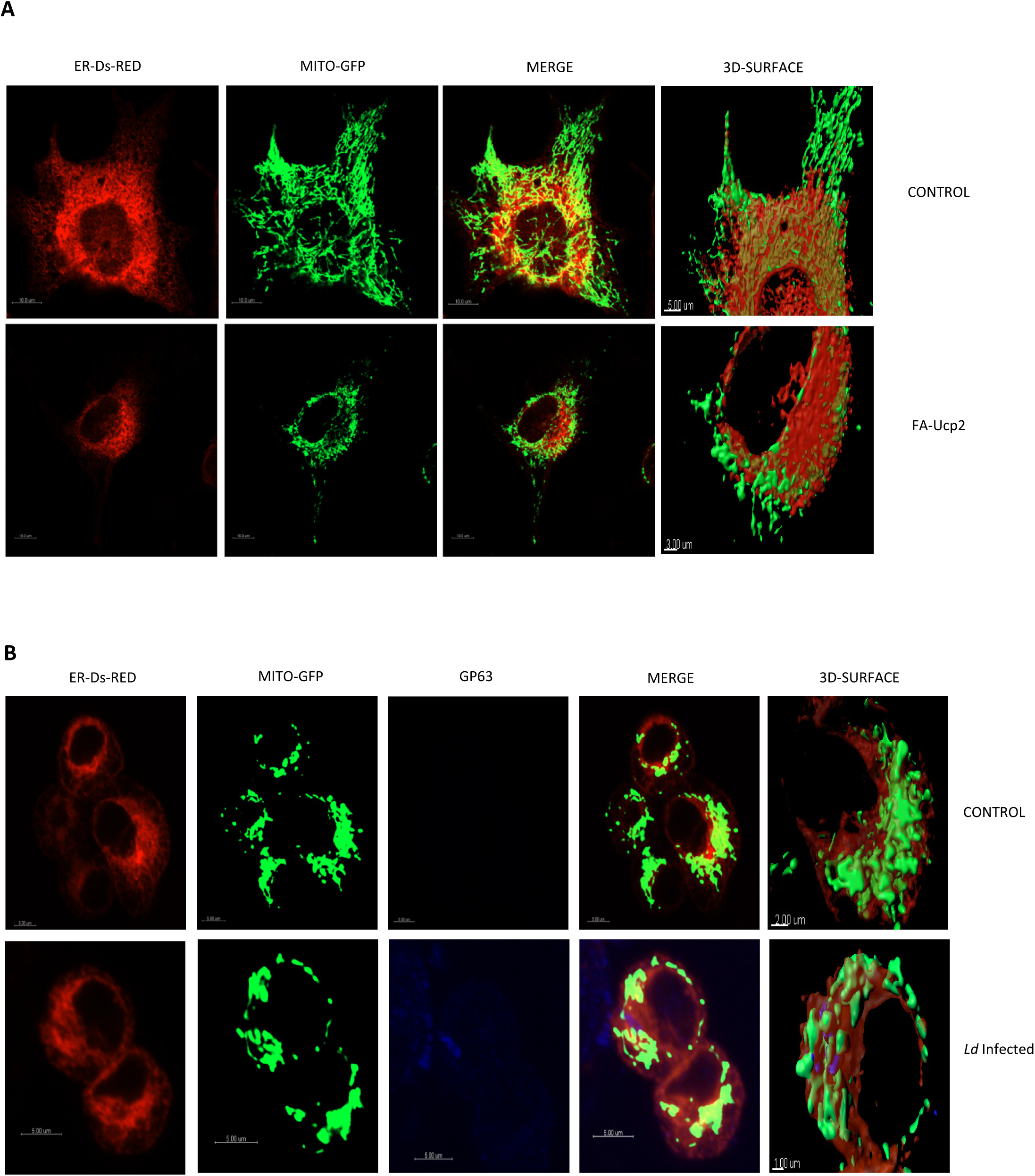
Effect of FH-Ucp2 expression and Leishmania infection on ER-mitochondria interaction in HeLa and RAW264.7 cells. (A) Microscopy of HeLa cells documenting the ER (depicted in red) and mitochondria (depicted in green) interaction in cells untransfected or transfected with FH-Ucp2. (B) Microscopy of RAW cells to show the ER (depicted in red) and mitochondria (depicted in green) interaction in cells uninfected or infected with *Leishmania*. The Leishmanial protein GP63 was depicted in blue to show infection.

**Supplemental Table S1.** Details of Plasmids and miRNA mimics.

| **Name of the Plasmid** | **Reference/Source** | **Plasmid Description** |
| --- | --- | --- |
| pmiR122 | As described by Ghosh et al 2013 (Ghosh et al., 2013) | Plasmid encoding pre-miR122 under a constitutive U6 promoter. |
| FLAG-HA-AGO2 | From Gunter Meister (Meister et al., 2004) | FLAG and HA tagged human AGO2 expression plasmid |
| NHA-LacZ | From Witold Filipowicz (Pillai et al., 2005) | NHA tagged version of LacZ protein |
| NHA-AGO2, NHA-AGO3 | From Witold Filipowicz ((Pillai et al., 2005) | N-HA tagged human AGO2, AGO3 expression plasmid |
| pRL-Con | From Witold Filipowicz (Pillai et al., 2005) | Humanized Renilla Luciferase (RL) coding region |
| pRL-Per-miR-122 | Described earlier (Basu and Bhattacharyya, 2014) | Perfect miR-122 binding site downstream of RL coding region |
| RL-Per-let7a | From Witold Filipowicz (Pillai et al., 2005) | Perfect let-7a binding site downstream of RL coding region |
| pRL-3Xbulge-miR-122 | From Witold Filipowicz (Bhattacharyya et al., 2006) | Three miR-122 binding sites downstream of Renilla Luciferase (RL) coding region. |
| PGL3-FF | From Promega | Firefly Luciferase (FL) under SV40 Promoter. |
| FLAG-HA-UCP2 | Described earlier (Chakrabarty and Bhattacharyya, 2017) | FLAG and HA tagged UCP2 expression plasmid |
| YFP-Endo | From Edouard Bertrand | YFP-tagged early endosome expression plasmid |
| CD63-GFP | From Edouard Bertrand | GFP tagged tetraspanin CD63 expression plasmid |
| ER-Ds-Red | From Clontech | Expression plasmid with a Ds Red Protein with ER targeting sequence |
| Mito-GFP | From Clontech | Expression plasmid with GFP Protein with mitochondria targeting sequence |
| pre-miR-122 mimic | From Life technologies (PM11012) |  |
| pre-miR-146a mimic | From Life technologies (PM10722) |  |

**Supplemental Table S2.**
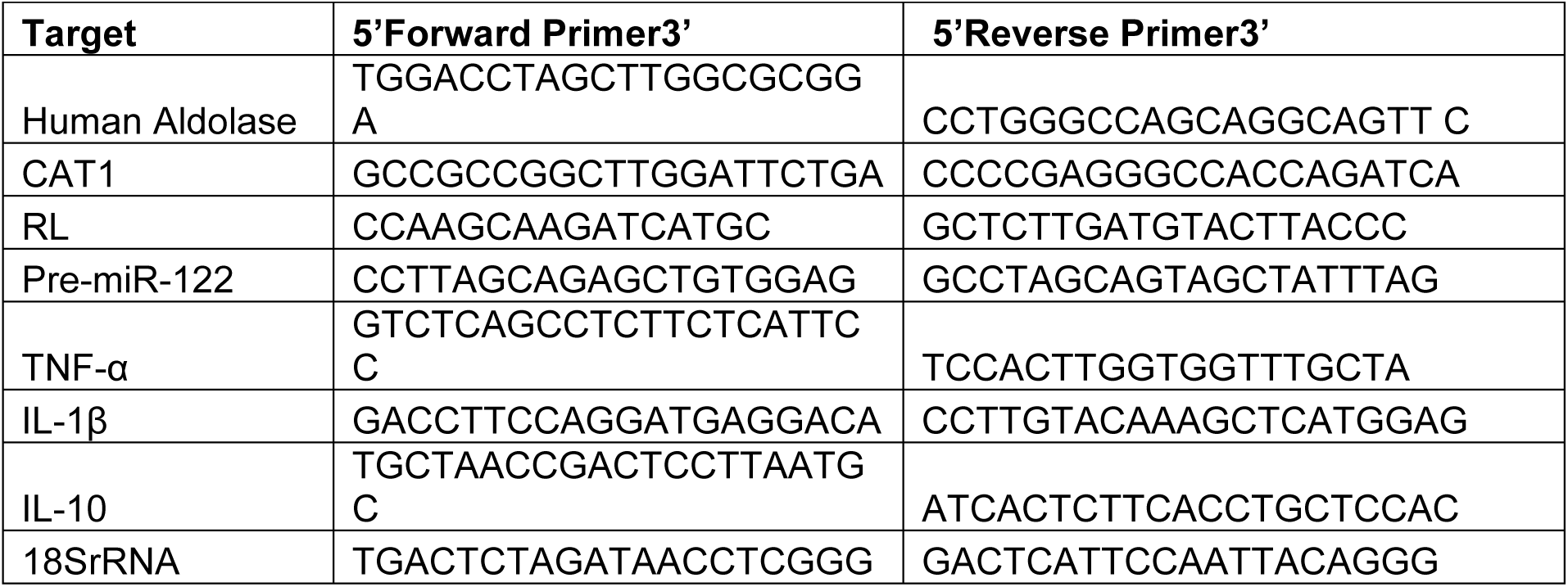
Detail of Primers used.

**Supplemental Table S3.** miRNA assays used for quantitative purpose.

| NAME | ASSAY ID |
| --- | --- |
| human miR-122 | assay ID 000445 |
| human miR-122* | assay ID 002130 |
| human let-7a | assay ID 000377 |
| human miR-16 | assay ID 000391 |
| mouse miR-155 | assay ID 002571 |
| human miR-146a | assay ID 000468 |
| U6 snRNA | assay ID 001973 |

**Supplemental Table S4.** Antibodies used.

| Antibody names | Raised in | Dilution | Source |
| --- | --- | --- | --- |
| AGO2 (EIF2C2) | Mouse | 1:1000 | Abnova |
| ALIX | Mouse | 1:200 | Santa cruz |
| β-Actin | Mouse Monoclonal (HRP conjugated) | 1 :10000 | Sigma Aldrich |
| HA | Rat monoclonal | 1:1000 | Roche |
| HRS | Rabbit | 1:1000 | Bethyl |
| Dicer1 | Rabbit | 1:5000 | Bethyl |
| CD63 | Mouse | 1:1000 | BD Pharmingen |
| Rab5 | Rabbit | 1:1000 | Cell Signaling |
| Rab7 | Rabbit | 1:1000 | Cell Signaling |
| RILP | Goat | 1:1000 | Santa cruz |
| UCP2 | Goat | 1:5000 | Novus |
| Dynamin2 | Rabbit | 1:1000 | Cell Signaling |
| Calnexin | Rabbit | 1:8000 | Bethyl |
| LAMP1 | Rabbit | 1:1000 | Cell Signaling |
| LC3b | Rabbit | 1:1000 | Cell Signaling |
| phospho-eIF2α | Rabbit | 1:1000 | Cell Signaling |
| eIF2α | Rabbit | 1:1000 | Cell Signaling |
| Rab27b | Rabbit | 1:1000 | Cell Signaling |

**Supplemental Table S5.**
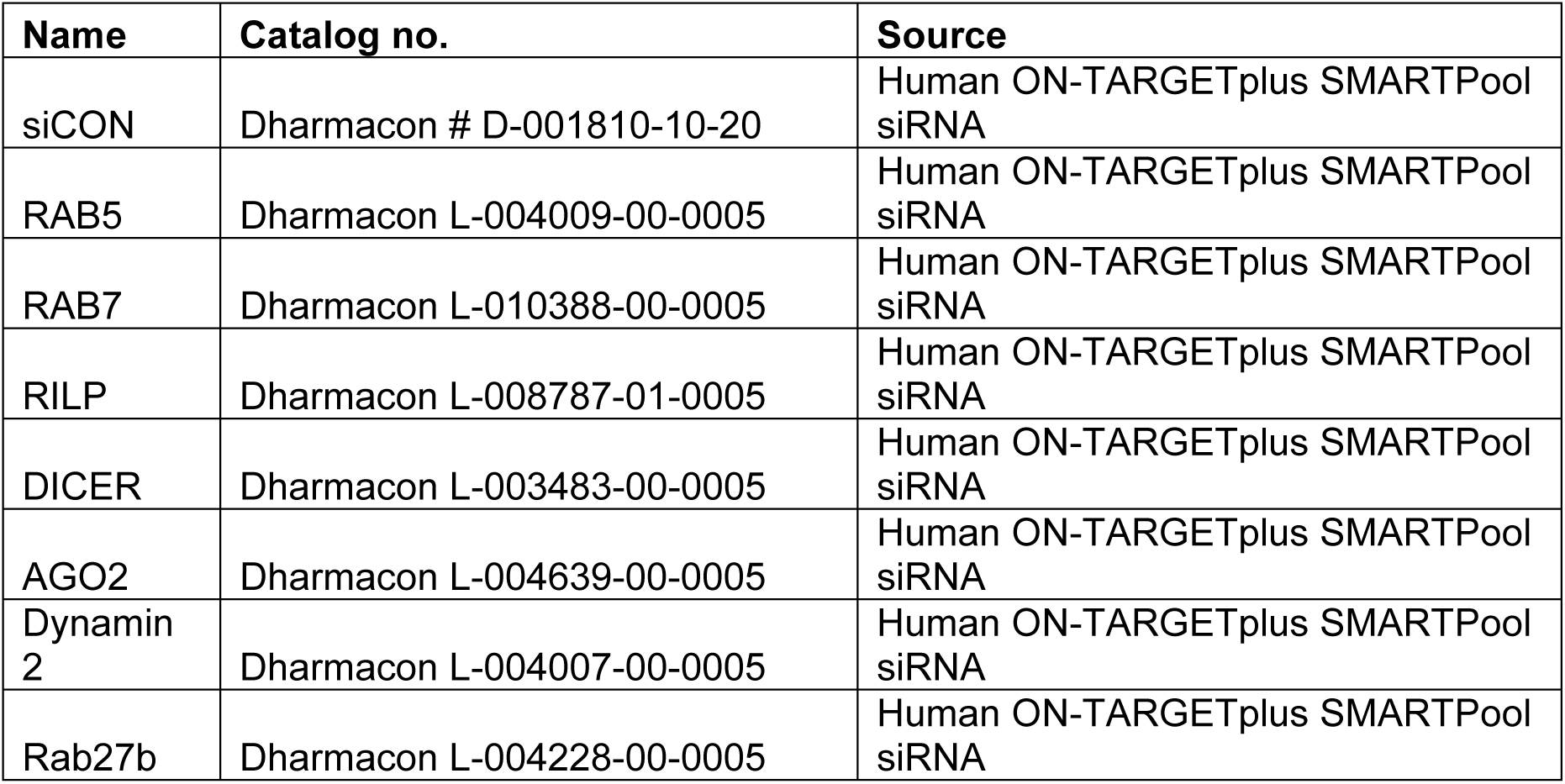
siRNAs Used.

## Notes

### Competing Interest Statement

The authors have declared no competing interest.

